# Spatiotemporal proteomics elucidates trafficking and localization of Neuroligins to the axon initial segment plasma membrane

**DOI:** 10.64898/2026.06.18.733123

**Authors:** Noortje Kersten, Chun Hei Li, Viktor al Naqib, Lukas de Haas, Victoria Vataka, Eva van der Meer, Peter Uljee, Guillaine Levenswaard, Maarten Altelaar, Amélie Fréal, Maarten Kole, Ginny G. Farías

## Abstract

Neuronal development and function rely on precise sorting of membrane proteins to distinct neuronal domains, yet the underlying mechanisms remain incompletely understood. Here, we investigated the trafficking of two highly homologous NLGNs (NLGN1 and 2) which distinctly localize and function at excitatory and inhibitory synapses. By using spatiotemporal proteomics, genetic engineering and microscopy in CNS cultured neurons and organotypic slices, we dissected the itinerary of biosynthetic NLGNs, identifying co-cargoes and sorting routes to dendrites and the AIS. At the AIS, which contains only inhibitory synapses, both NLGN1 and NLGN2 are locally exocytosed. While NLGN2 is stably recruited to axo-axonic synapses by its extracellular domain and modulated by activity, NLGN1 is retrieved by endocytic mechanisms. We propose that biosynthetic NLGNs are co-sorted to the AIS PM where they detect pre-synaptic type. These findings may help to elucidate the role of AIS-localized NLGNs in shaping AIS structural and functional plasticity.

**Highlights:**

- Spatiotemporal proteomics elucidates interactome of biosynthetic NLGN1 in neurons
- NLGN1 and NLGN2 are targeted to the AIS by kinesin-1
- NLGN2, but not NLGN1, is retained at axo-axonic synapses and modulated by activity
- NLGN stability at the AIS plasma membrane is determined by the ectodomain

**Graphical Abstract:** 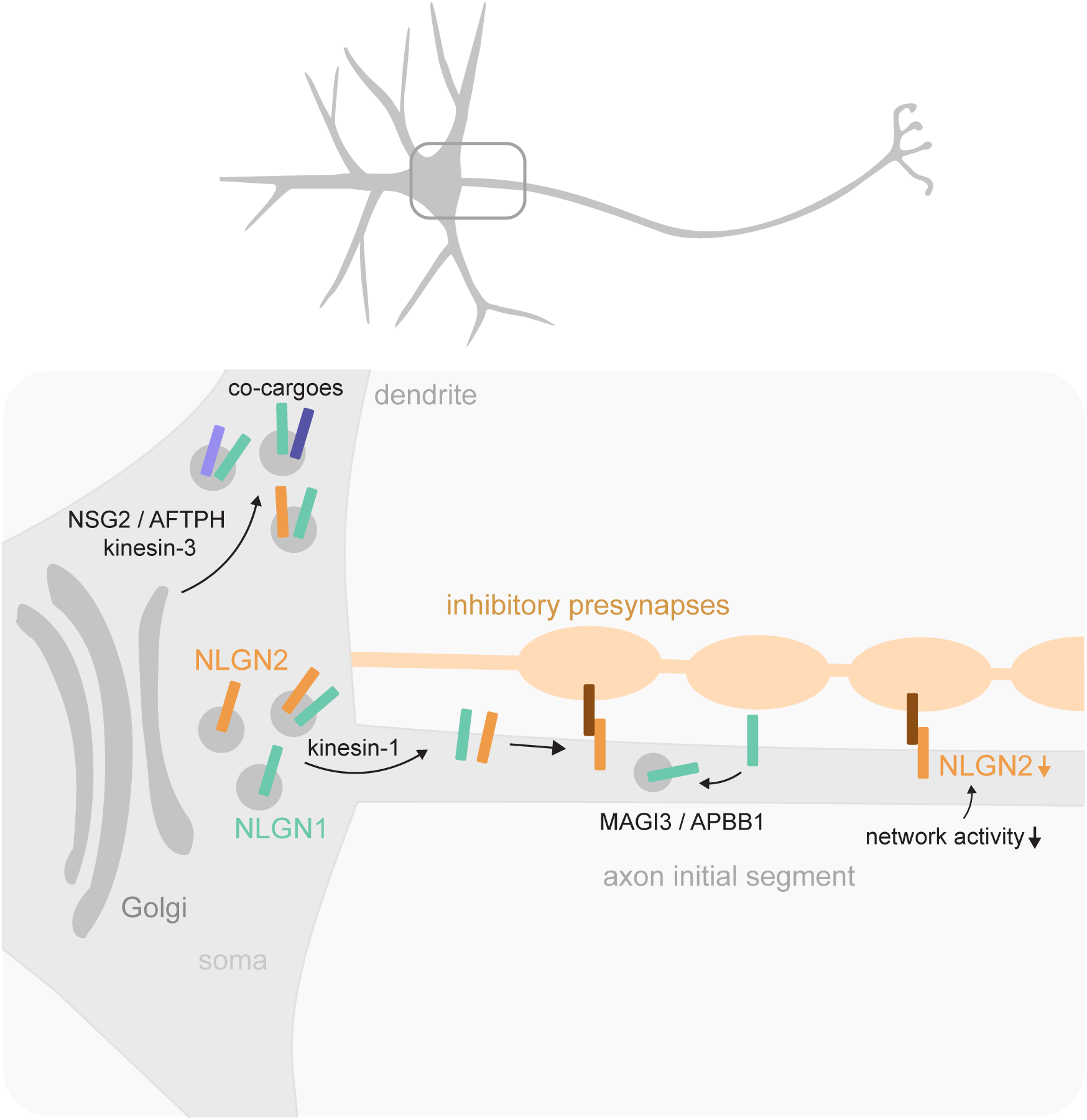

## Introduction

Neurons are highly polarized cells that must maintain a unique proteome in dendrite and axon subdomains for proper neuronal development and function. Central to this organization is the correct sorting and trafficking of newly synthesized proteins from the Golgi apparatus located in the neuronal soma. Foundational studies have revealed the main characters orchestrating this: Adaptor Protein (AP) complexes recognize and sort biosynthetic proteins (cargoes) into specific vesicle types at the Trans-Golgi-Network (TGN). For example, AP-1 and AP-4 sort dendritic cargoes^1–3^, while AP-3 sorts axonal cargoes into vesicles for the delivery to their respective neuronal domains^3^. Then, specific MT-driven motor proteins deliver distinct vesicles to the correct destination^4–6^. Yet, emerging research is revealing that this system is more intricate than it was once thought. For instance, only few dendritic and axonal biosynthetic cargoes have the same classical motifs to be recognized by either AP-1, AP-3 or AP-4 at the TGN, and many other cargoes do not contain such motifs^4,6^. In addition, some cargo families, like Neuroligins (NLGNs), with high homology in their cytosolic tails need to be delivered to distinct subdomains.

The Neuroligin (NLGN1-4) family represents a prime example of highly homologous cargoes that lack canonical motifs for adaptor protein recognition during TGN-to-dendrite sorting. NLGNs are somatodendritic synaptic adhesion molecules, which play essential roles in synaptic maturation and function^7–9^. Their dysfunctions are linked to neurodevelopmental and neuropsychiatric disorders (e.g. autism, epilepsy, and schizophrenia)^10–12^. Mutations in NLGNs, primarily found in their luminal/extracellular domain, are structurally destabilizing, suggesting they impair NLGN sorting within the biosynthetic pathway^13^.

NLGNs distribute to specific dendritic synaptic subdomains. For instance, NLGN1 is enriched at glutamatergic excitatory synapses, while NLGN2 is enriched at GABAergic inhibitory synapses, and NLGN3 is present at both synapses^7^. The correct localization of different NLGNs maintains excitation/inhibition balance^14,15^. Despite their different locations and functions, NLGNs biochemically interact with the same extra- and intracellular partners (e.g. neurexins and gephyrin, respectively)^16^. Extending the complexity of this puzzle, a recent study found that biosynthetic NLGN1 is transiently sorted to the axon initial segment (AIS) plasma membrane (PM)^17^; however, neither the mechanism through which this happens, nor the functional relevance has been elucidated. Overall, little is known about the mechanisms governing biosynthetic NLGN sorting to the correct neuronal subdomain for their local functions at synapses.

Identifying the sorting mechanisms of biosynthetic cargo is challenging due to their steady-state distribution at destination sites and the technical limitations of analyzing only a few interactors at a time. We recently developed POTATOMap (Protein Origin, Trafficking And Targeting to Organelle Mapping) tool for the spatiotemporal profiling of biosynthetic cargo and its transient interactome through proximity labeling and proteomics^18^. This tool allowed the identification of co-cargoes and the mechanisms involved in the sorting of biosynthetic lysosomal proteins to lysosomes located at different neuronal domains^18^.

Here, we applied POTATOMap to understand sorting mechanisms of NLGNs in CNS neurons. Using biosynthetic NLGN1 as a bait, we identified a subset of dendritic proteins that co-traffic with NLGN1, and adaptor and motor proteins involved in their polarized sorting towards dendrites. Moreover, we uncovered trafficking of NLGN1 and NLGN2 to the AIS, which is mediated by kinesin-1. We found that NLGN2, but not NLGN1, is part of inhibitory axo-axonic synapses at the AIS, in cultured neurons and organotypic slice cultures (OSCs). Most NLGN1 at the AIS is removed by endocytic mechanisms, while NLGN2 is retained and regulated by neuronal activity. We propose a model in which NLGNs are trafficked together, and only upon recognition of respective presynaptic partners, a selective retention mechanism segregates the different paralogs into excitatory versus inhibitory synapses. Our results open new avenues to explore how AIS-localized NLGNs shape AIS structural and functional plasticity.

## Results

### Spatiotemporal interactome of biosynthetic NLGN1 during its trafficking from ER to PM

Comprehensive data on polarized trafficking in neurons is lacking. To address this gap, we applied the POTATOMap tool for spatiotemporal mass spectrometry of secretory cargoes^18^, to the somatodendritic postsynaptic adhesion protein NLGN1 (**Fig. 1A**). This tool combines the Retention Using Selective Hooks (RUSH) and the APEX2 systems^19,20^ to gain spatiotemporal control over biosynthetic NLGN1 and determine its interactome over time (**Fig. 1A**). Briefly, newly synthesized NLGN1 is tagged with luminal Streptavidin Binding Peptide (SBP) for the RUSH system, a cytoplasmic V5 tag for cargo tracking, and APEX2 for proximity labeling and proteomics (RUSH-NLGN1-V5-APEX2). Its co-expression with Streptavidin (Strep) fused with a KDEL ER retention signal causes Strep-SBP heterodimerization, efficiently retaining biosynthetic NLGN1 at the ER immediately after synthesis (**Fig. 1A, B**). The addition of RUSH and APEX2 modules to NLGN1 does not alter its sorting as shown by its co-distribution with PSD95 at dendritic excitatory synapses, in the absence of the Strep-KDEL retention system (**Fig. 1C**). Upon addition of biotin, which outcompetes SBP for binding to Strep-KDEL, NLGN1 is released from the ER and its trafficking through the biosynthetic pathway is synchronized (**Fig. 1A, D**).

**Figure 1.**
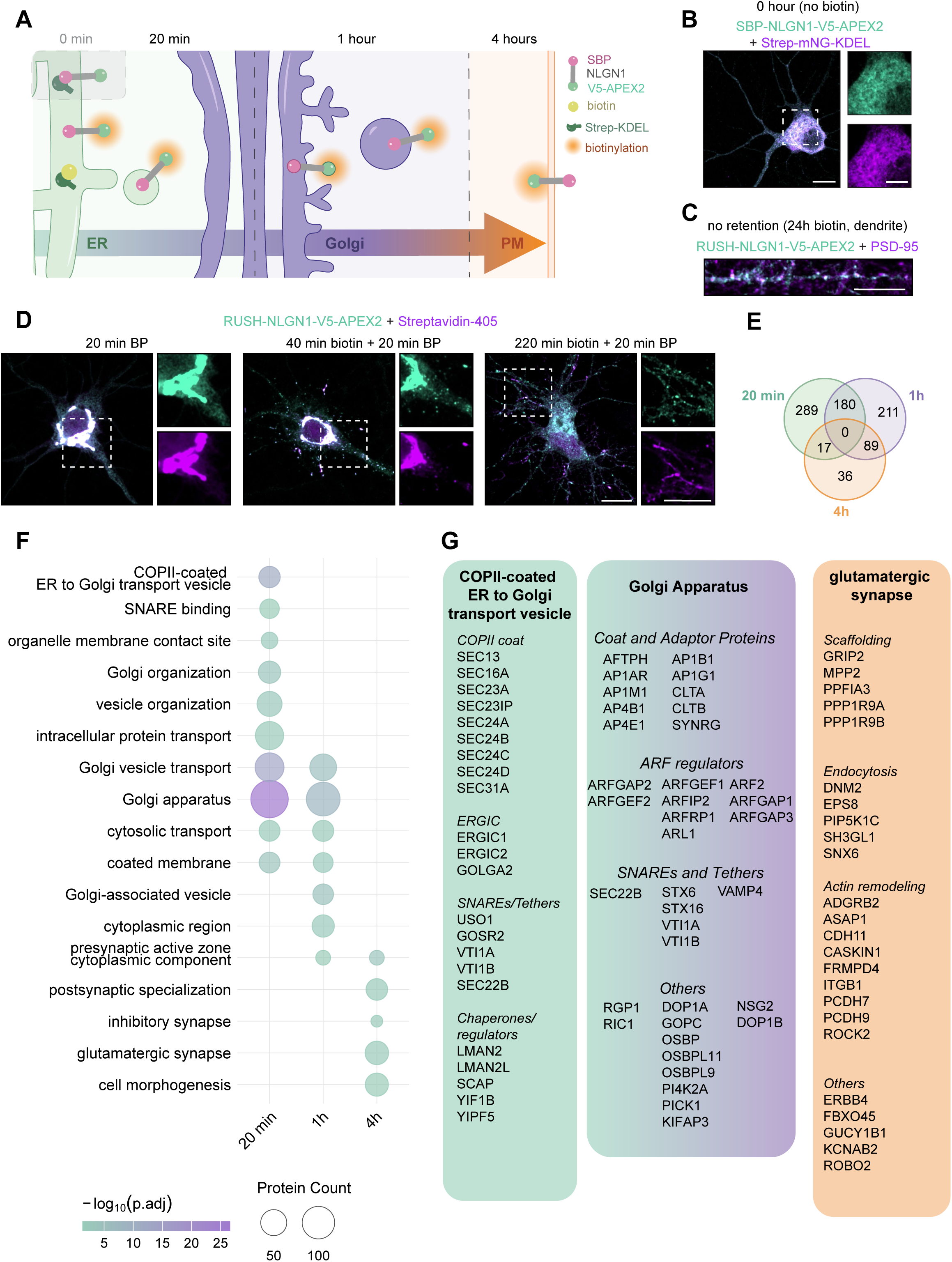
Spatiotemporal interactome of biosynthetic NLGN1 during its trafficking from ER to PM. (A) Schematic of the POTATOMap system, applied to NLGN1. (B) Representative image of a DIV8 cortical neuron tranduced with lentivirus containing SBP-NLGN1-V5-APEX2 and Strep-mNG-KDEL vectors, cultured without biotin to retain the cargo in the ER. (C) Representative image of a dendrite showing normal distribution of RUSH-NLGN1-V5-APEX2 at excitatory synapses in neurons cultured with biotin to prevent ER retention and stained for V5 and PSD-95. (D) Representative images of neurons transduced with lentiviruses for SBP-NLGN1-V5-APEX2 and Strep-mNG-KDEL. Neurons were incubated with biotin and BP as indicated, and 1 min. H2O2. Cells were stained for V5 and for biotin with Streptavidin-405. (E-G) Venn diagram of proteins significantly enriched at their respective time point(s). N = 2 independent experiments with 2 technical repeats per experiment; missing values were imputed for statistical purposes. ANOVA with Tukey’s correction, *p* value <0.05, appropriate log_2_ fold change thresholding was set per time point (E). Gene Ontology enrichment analysis of the proteins in each time point, *p* value <0.05 (adjusted with Benjamini-Hochberg FDR correction (F). Schematic representation of proteins within enriched terms. Within these terms, proteins were further segregated manually according to known function. For the Golgi Apparatus, we depicted a selection of proteins (G). Scale bars in overview images represent 10 µm, scale bars in zooms represent 5 µm. See also Fig. S1.

We characterized the trafficking dynamics of RUSH-NLGN1-V5-APEX2 in neurons at day-in-vitro (DIV)-8 and established three time points that reveal distinct steps in NLGN1’s biosynthetic pathway. After 20 min., NLGN1 is moving from the ER to the Golgi. After 1h, we observed NLGN1 signal at the Golgi and in vesicles. After 4h, NLGN1 has reached its destination in the dendrites (**Fig. 1D, S1A**). These dynamics also become apparent in western-blots of NLGN1 at these time points, where a clear shift from immature to mature glycosylated NLGN1 can be observed, indicating processing at the Golgi (**Fig. S1B**). To activate APEX2, we incubated the cells with biotin-phenol for 20 minutes in the presence or absence of H_2_O_2_ for 1 min. at the end of each time point (**Fig. 1D, S1B**). Biotinylated proteins were then isolated through streptavidin-pull down and analyzed by LC-MS/MS. All hits were first tested against the -H_2_O_2_ control and then between time points.

POTATOMap detected a total of 2353 proteins. Testing between time points revealed an enrichment of 486 proteins at 20 min., 480 proteins at 1h and 142 proteins at 4h (**Fig. 1E**). GO term analysis of these differentially enriched proteins further illustrates NLGN trafficking through the biosynthetic pathway. At the 20 min. point, terms related to ER-Golgi transport were enriched. At the 1h point, terms related to post-Golgi trafficking were enriched. Finally, at the 4h time point, various synaptic terms were enriched (**Fig. 1F**). We further highlighted three of these enriched terms (**Fig. 1G**). We identified components of the COPII vesicle coat, as well as ERGIC components during NLGN1 sorting along the early secretory pathway (**Fig. 1G**). Later, we observed subunits of the Adaptor Protein complexes, several ARF regulators, SNAREs, and the known interactor GOPC (**Fig. 1G**). Finally, we detected several components enriched at glutamatergic synapses, such as endocytosis and actin remodeling machineries (**Fig. 1G**). Some proteins were detected at just one time point; these are referred to as ‘unique proteins’. We performed a STRING Gene Enrichment analysis on the unique proteins for each time point (**Fig. S1C**).

To test the specificity of the tool for secretory pathways, we compared our biosynthetic NLGN1 interactome with our previously established biosynthetic lysosomal protein LAMP1 interactome. We found that most proteins were shared between NLGN1’s 20 min. point and LAMP1’s 1h point, indicating different trafficking dynamics of these two secretory cargoes (**Fig. S1D**). STRING Gene Enrichment analysis revealed that most of these shared proteins are components of the secretory pathway (**Fig S1E**). The vast majority of identified proteins, however, were unique to either cargo, revealing their differential sorting routes. This illustrates the specificity of POTATOMap and further confirms it as a powerful tool to study distinct biosynthetic pathways.

### NLGN1 is co-trafficked with a subset of secreted cargoes

Next, we utilized our NLGN1 POTATOMap MS data to define the vesicle population(s) in which biosynthetic NLGN1 is trafficked, identifying co-cargoes.

For this, the proteins from all trafficking time points (20 min. and 1h time points) were pulled and filtered for membrane proteins, showing a total of 269 proteins. From these 269 membrane proteins, STRING Gene Enrichment analysis showed that 113 proteins are sorted to the PM, from now on referred to as co-cargoes (**Fig. 2A**). Most of these proteins decreased or were constant in their abundance over time, indicating that they segregate, or stay in close proximity to NLGN1 (**Fig. 2B**). Interestingly, a comparison of these 113 proteins with all 227 postsynaptic PM proteins revealed that merely a small group of 55 postsynaptic membrane proteins potentially traffic together with NLGN1 (**Fig. S2A**). This could mean that postsynaptic membrane proteins are trafficked to the synapse in distinct vesicle populations, segregated by place, time, or both. We plotted the temporal signature of a selection of these co-cargo candidates displaying very minor changes in abundance over time (**Fig. 2C**). Interestingly, within these, we observed the NLGN1 paralogs NLGN2 and NLGN3 as co-cargoes (**Fig. 2C**), suggesting they share same secretory pathway.

**Figure 2.**
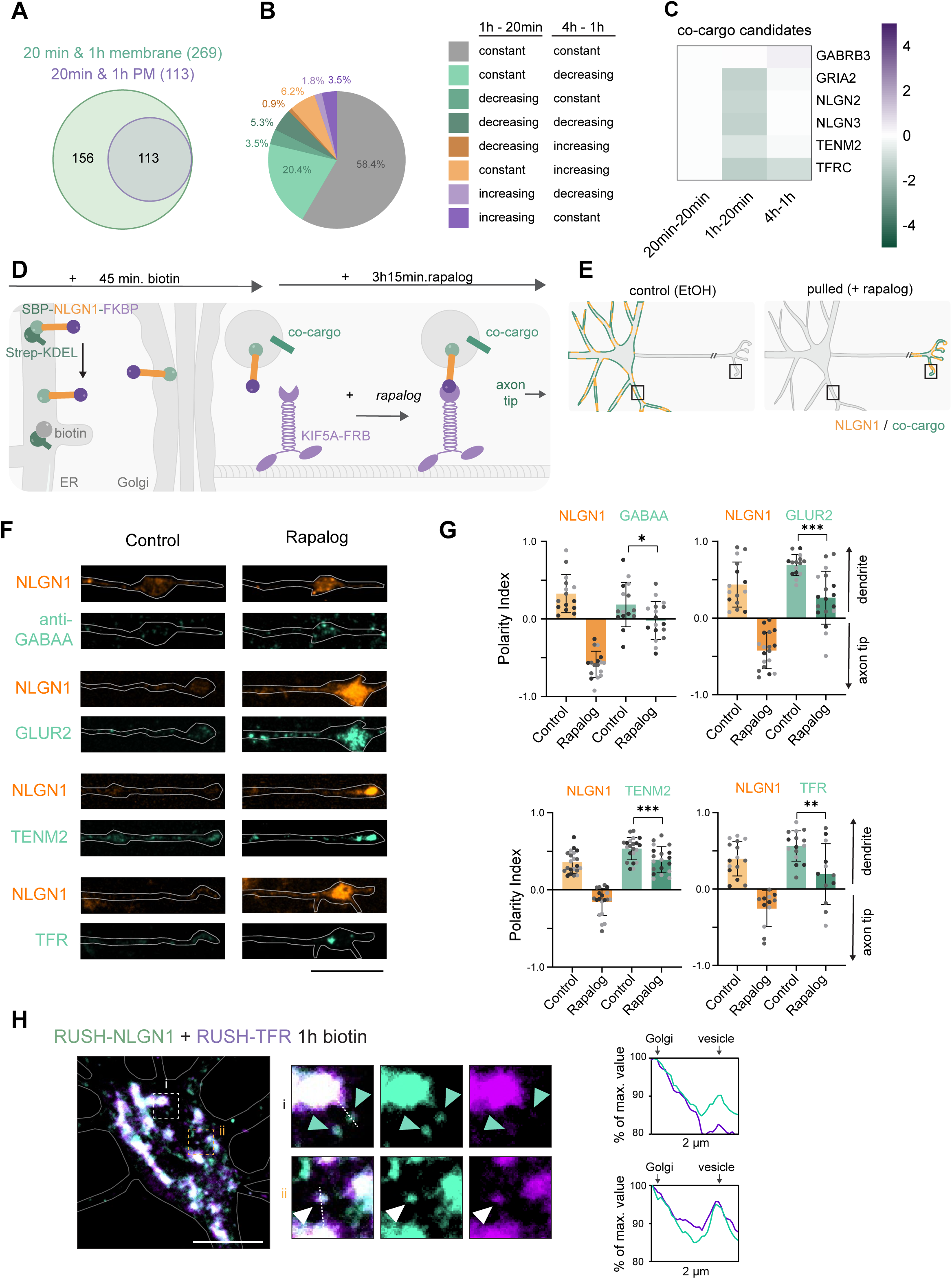
NLGN1 co-traffics with a subset of secreted cargoes. (A-C) Venn diagram of 20 min. & 1h proteins with a transmembrane domain, and 20min. & 1h transmembrane proteins annotated to the PM, termed co-cargoes (A). Pie chart showing the temporal profile of 20min & 1h plasma membrane proteins (B). Heat map of selected co-cargo candidates in which the log2 Fold Change across time points is plotted (C). (D-E) Schematic representation of the axonal relocalization system (D) and of the expected outcome if a candidate co-cargo is trafficked together with NLGN1 (E). (F-G) Representative images of axon tip of DIV8 hippocampal neurons transfected with SBP-NLGN1-Halo-FKBP and HA-KIF5A-FRB or mCherry-KIF5A-FRB (not shown), stained for GABAA, or co-transfected with either HA-GLUR2 or TFR-GFP or TENM2-EGFP, treated as shown in (D), with biotin and with EtOH (control) or Rapalog (F). Quantification of Polarity Index. Only cells with relocalized NLGN1 were included. (G). Each dot represents a neuron, each color an independent experiment. Polarity Index for GLUR2 and TFR were analyzed with unpaired t test with Welch’s correction, and GABAA and TENM2 with unpaired t test. *p < 0.05, **p < 0.01, and ***p < 0.001, ****p < 0.0001. (H) Representative still image of a live neuron expressing RUSH-NLGN1-mNG and Halo-TFR-RUSH, incubated with biotin for 1h. Zoom and intensity profile lines (left). Arrows indicate vesicles positive for just NLGN1 (green) or both NLGN1 and TFR (white). Scale bars represent 10 µm. See also Fig. S2.

Next, we verified the presence of potential co-cargoes on the same post-Golgi vesicles as NLGN1. We established a tool to relocate and enrich biosynthetic NLGN1-positive carriers into the axon tip. To this end, we appended a Halo-FKBP module on the cytosolic side of our RUSH-NLGN1 construct, while co-expressing the motor domain of the axonal motor KIF5A, tagged with FRB. After 45 min. biotin, RUSH-NLGN1 vesicles are budding from the Golgi (**Fig. 2D**). At this point, Rapalog was added to induce FKBP/FRB dimerization, which couples the post-Golgi NLGN1-carriers to the axonal motor to drive them into the axon tip (**Fig. 2D, E**). Importantly, this did not result in Golgi fragmentation or relocation of Golgi components to the axon (**Fig. S2B, C**). Using this vesicle relocation tool, we assessed whether the potential co-cargoes are also relocated to the axon tip, thereby validating their presence on the same carrier. Relocalization was assessed by quantifying the Polarity Index between dendrites and the axon tip (see Methods). Using this approach, we were able to validate GABAA, GLUR2/GRIA2, TENM2 and TFR as co-cargo of NLGN1. After successful relocalization of NLGN1 to the axon tip, these cargoes were significantly redistributed from the dendrites to the axon tip (**Fig. 2F, G**). Important to note, the studied co-cargoes were analyzed at the steady state, meaning that just a small pool corresponded to biosynthetic cargo, which we could still visualize co-trafficked with biosynthetic NLGN1 to the axon tip.

While the axonal relocalization assay tells us that multiple cargoes are trafficked together with NLGN1, it does not inform us if co-cargoes are always on the same biosynthetic vesicle, or whether there are vesicle subpopulations. To investigate this, we performed live-cell imaging of biosynthetic NLGN1 and biosynthetic TFR using the RUSH system. We observed vesicles in the soma that were positive for both NLGN1 and TFR, as well as vesicles that were positive for only NLGN1 (**Fig. 2H, S2D**). This suggests that NLGN1 can exit the Golgi in different vesicle subpopulations.

Thus, we find an entire set of cargoes transported along the same route as biosynthetic NLGN1, shedding light on the existence of vesicle subpopulations.

### Dendritic sorting and trafficking of NLGN1 is mediated by AFTPH, NSG2, and kinesin-3

Next, we aimed to elucidate the sorting machineries of biosynthetic NLGN1. In our POTATOMap dataset, we identified 3 out of 4 subunits of the AP-1 complex. These were most enriched at the 20 min. time point and were declining towards the 1h and 4h time points (**Fig. 1G**; **Fig. 3A**). We also found 2 out of 4 subunits of the AP-4 complex to be uniquely present at the 20 min.-mark (**Fig. S1D**) and 2 subunits of the AP-3 complex (**Fig. 3A**). AP-1 and AP-4 are known to sort cargo from the Golgi into the dendrites. A known dendritic cargo of AP-1 is TFR, whereas a known cargo of AP-4 is the AMPA receptor GLUR2 ^1,2^. The fact that both TFR and AMPA receptor co-traffic with NLGN1 (**Fig. 2**), suggests that NLGN1 is a promiscuous cargo that can be sorted by either AP complex.

**Figure 3.**
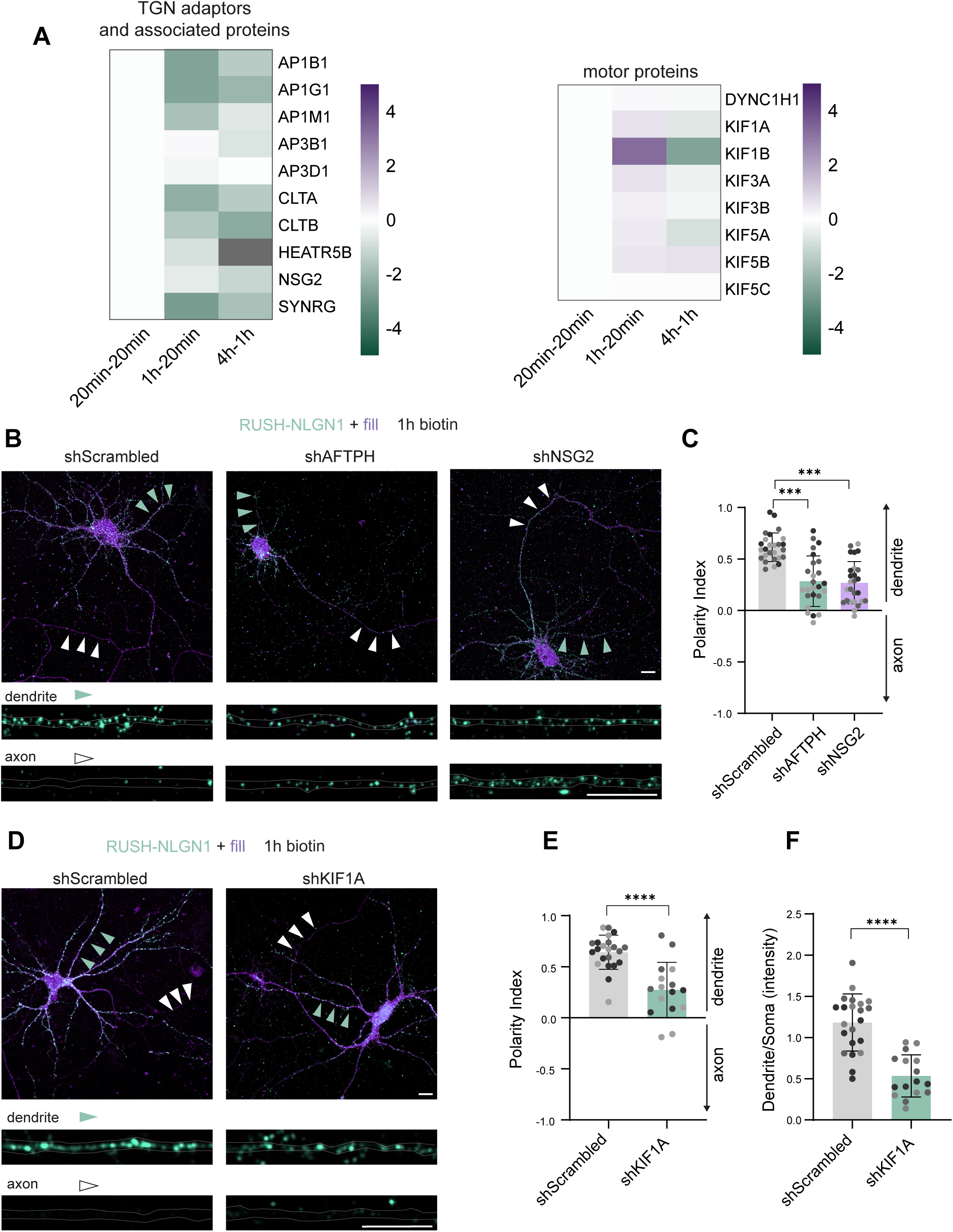
Dendritic sorting and trafficking of NLGN1 is mediated by AFTPH, NSG2, and kinesin-3. (A) Heat map of TGN adaptors and associated proteins (left) and motor proteins (right) in which the log2 Fold Change across time points is plotted. (B-C) Representative images of neurons expressing RUSH-HA-NLGN1, tdTomato (fill) and shScrambled, shAFTPH or shNSG2, live incubated with biotin for 1h and surface labeled for HA (B). Quantification of Polarity Index (C). (D-F) Representative images of neurons expressing RUSH-HA-NLGN1, tdTomato (fill) and shScrambled or shKIF1A, live incubated with biotin for 1h and surface labeled for HA (D). Quantification of Polarity Index (E). Quantification of Dendrite/Soma intensity ratio (F). Green arrows indicate dendrites and white arrows the axon. Data points represent individual neurons, color indicates individual experiments. Brown-Forsythe & Welch ANOVA with Dunnett T3 correction (C), unpaired t test with Welch’s correction (E), unpaired t test (F). *p < 0.05, **p < 0.01, ***p < 0.001, ****p < 0.0001. Scale bars represent 10 µm. See also Fig. S3.

Interestingly, we also found two less well-known proteins possibly involved in NLGN1 biosynthetic sorting at the TGN: Aftiphilin (AFTPH) (**Fig. S1D**) and Neuron Specific Gene-2 (NSG2) (**Fig. 3A**). AFTPH is a brain-enriched protein, localizing to the TGN and clathrin-coated vesicles. It was identified as a binding partner of AP-1 and AP-2 in *in vitro* assays and brain extracts^21,22^. NSG2/P19 is a membrane protein, localized to the TGN^23^, and by similarity to NSG3, it is predicted to bind clathrin^24^. To study the role of AFTPH and NSG2 in the sorting of biosynthetic NLGN1 to the dendritic PM, we first made a RUSH-NLGN1 with an N-terminal (extracellular) HA tag (RUSH-HA-NLGN1). We released RUSH-NLGN1 from the ER for 1h, while labeling the surface to detect its delivery to the PM, followed by secondary labeling after fixation prior to permeabilization. The surface HA labeling only detects a pool of the total NLGN1 that has reached the dendritic surface, as shown with labeling of an mNG-and HA-tagged RUSH-NLGN1 construct (**Fig. S3A**). The surface distribution of NLGN1 was quantified by measuring the Polarity Index between dendrite and the axon for neurons expressing scrambled shRNA control, or shRNAs targeting AFTPH or NSG2. Knockdown of AFTPH and NSG2 impaired the distribution of biosynthetic RUSH-NLGN1 to dendrites, after 1h release (**Fig. 3B, C; Fig. S3B, C**). This indicates that AFTPH and NSG2 contribute to the polarized sorting of NLGN1-positive vesicles to dendrites.

In addition, in our POTATOMap MS data, we identified different kinesin motors and components of the dynein complex (**Fig. 3A**), as well as their adaptors (**Fig. S3D**). Interestingly, we noticed the transient enrichment of kinesin-3 (KIF1A and KIF1B) at 1h and its reduction at 4h (**Fig. 3A**). Kinesin-3 has been shown to transport dense core vesicles (DCVs) into the axon and dendrites^25,26^. Moreover, we also detected the scaffolding proteins Liprin-α (PPFIA) and TANC2, which are known to subsequently recruit the KIF1A-DCV complexes into dendritic spines (**Fig. S3D**)^26^. It could therefore be possible that NLGN1 is transported to synapses by a similar mechanism as employed by DCVs. Therefore, we knocked down KIF1A and examined the sorting of RUSH-NLGN1 after 1h release (**Fig. 3D**). The knockdown of KIF1A caused reduced sorting of biosynthetic NLGN1 to dendritic PM. Instead, more NLGN1 was observed at the soma surface (**Fig. 3D-F**).

In summary, we identify key interactors of NLGN1 that mediate its sorting and trafficking to dendrites. These include AFTPH and NSG2 for selective sorting into dendritic vesicles at the TGN and coupling to kinesin-3 for vesicle transport to dendrites.

### NLGN1 and NLGN2 are transported to the AIS via kinesin-1

Interestingly, our POTATOMap data also revealed a transient enrichment of AIS-enriched proteins (e.g. ANKG/ANK3 and TRIM46), suggesting that biosynthetic NLGN1 visits the AIS during its trafficking (**Fig. 4A**). This is in line with a previous study, which described the transient sorting of biosynthetic NLGN1 to the AIS PM^17^.

**Figure 4.**
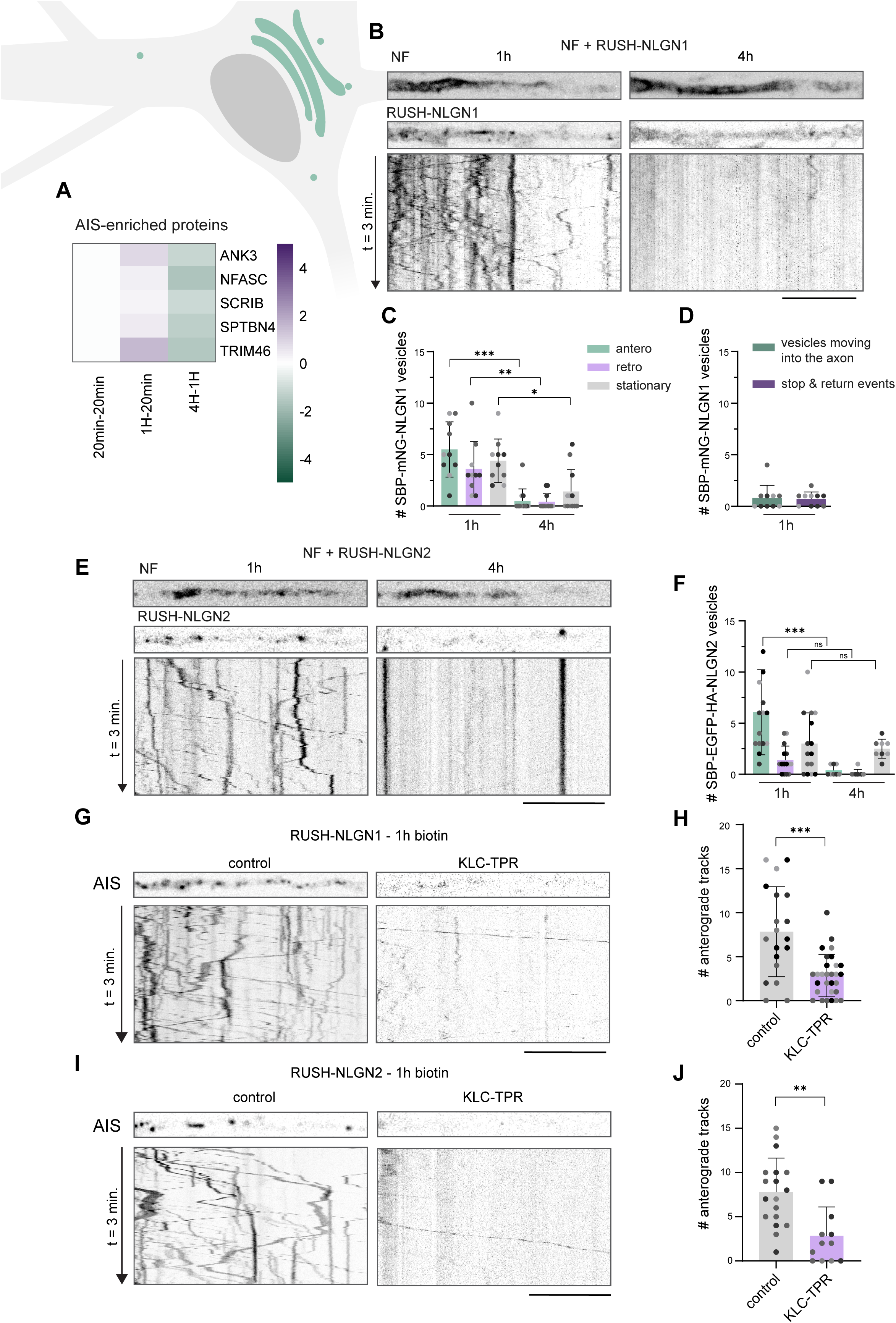
NLGN1 and NLGN2 are transported to the AIS by kinesin-1. (A) Heat map of AIS-enriched proteins, in which the log2 Fold Change across time points is plotted. (B-F) Still images and kymographs of the AIS of neurons expressing RUSH-mNG-NLGN1 (B) or RUSH-EGFP-HA-NLGN2 (E), incubated with biotin for 1h, then live imaged for 3 minutes, 1sec/frame. Quantification of NLGN1 (C) and NLGN2 (F) anterograde, retrograde and stationary tracks. Quantification of the number of NLGN1+ vesicles entering into the axon or stopping and returning towards the soma (D). (G-J) Still images and kymographs of the AIS of neurons expressing RUSH-mNG-NLGN1 (G) or RUSH-EGFP-HA-NLGN2 (I) with or without KLC-TPR. Quantification of the number of anterograde NLGN1 (H) or NLGN2 (J) tracks. Data points represent individual neurons, color represents individual experiments. Kruskal-Wallis test with Dunn’s correction (C), Mann-Whitney test (H), unpaired t-test (J). *p < 0.05, **p < 0.01, ***p < 0.001. Scale bars represent 10 µm. See also Fig. S4.

Here, we further characterized the movement of biosynthetic NLGN1 to the AIS by live cell imaging. For this, we labeled the AIS with Neurofascin (NF)-640 conjugated antibody prior to imaging and tracked the sorting of RUSH-NLGN1-positive vesicles for 3 min, after 1h and 4h biotin addition (**Fig. 4B**). From timelapse images, we generated kymographs to quantify the number of anterograde, retrograde and stationary vesicles at the AIS. Biosynthetic NLGN1 at 1h post-release showed a similar number of vesicles moving anterograde and retrograde, as well as stationary puncta at the AIS (**Fig. 4B, C**). However, after 4h, moving vesicles and stationary puncta have reduced significantly (**Fig. 4B, C**). This is in agreement with our POTATOMap data, indicating its transient visit to the AIS. Only a few vesicles were observed moving past the AIS into the axon after 1h release, suggesting this targeted sorting of NLGN1 is not due to overexpression artifacts (**Fig. 4D**). Moreover, we rarely observed NLGN1-positive vesicles stopping and immediately returning to the soma, indicative of the reported physical barrier and retrograde transport for other dendritic cargoes at the AIS^27–30^ (**Fig. 4D**).

In our spatiotemporal MS analysis, we identified several co-cargoes, including TFR, AMPA receptors, and NLGN1 paralogs (**Fig. 2C**). We wondered if this sorting route to the AIS is also employed by NLGN1 co-cargoes. However, we did not observe RUSH-TFR sorting to the AIS after 1h release (**Fig. S4A**). This aligns with previous studies indicating that neither biosynthetic TFR nor AMPA receptors are trafficked to the AIS^17,31^. Because of the high homology between NLGNs^32^, we then assessed whether biosynthetic NLGN2 is also trafficked into the AIS. Indeed, we observed a similar number of RUSH-NLGN2-positive vesicles entering the AIS after 1h biotin, and the number of anterograde tracks was reduced after 4h biotin. Overall, we observed less retrograde tracks than for NLGN1, and there were more stationary puncta at the 4h time point (**Fig. 4E, F**), suggesting a more stable distribution of NLGN2 at the AIS.

Next, we aimed to elucidate how NLGNs are transported to the AIS. We focused on the well-established axonal motors kinesin-1 (KIF5A-C) and kinesin-3 (KIF1A)^33^, which were identified in our proteomics dataset (**Fig. 3A**). We previously investigated the role of kinesin-3/KIF1A in dendritic transport (**Fig. 3D-F**). KIF1A knock-down, however, did not show any effect on biosynthetic NLGN1 trafficking to the AIS (**Fig. S4B, C**). To assess the involvement of kinesin-1, we expressed a dominant negative for kinesin light chain (KLC), KLC-TPR^34^, to inhibit the coupling of KIF5A-C to cargoes. Expression of the KLC-TPR construct significantly abolished anterograde trafficking of both NLGN1 and NLGN2 to the AIS (**Fig. 4G-J**).

In conclusion, we show that NLGNs are trafficked into the AIS by kinesin-1, but not kinesin-3, suggesting motor specificity for their sorting to the AIS and dendrites.

### While NLGN2 is stably inserted into the AIS PM, NLGN1 is removed by APBB1- and MAGI3-mediated endocytosis

Bourke et al. (2021) showed that biosynthetic NLGN1 is transiently inserted at the AIS PM by using a surface GFP antibody crosslinking protocol that prevents endocytosis or lateral diffusion of secreted cargoes. We confirmed this in our system by adapting their approach. RUSH-EGFP-NLGN1 was released from the ER for 2h, and during this period neurons were simultaneously incubated with a GFP polyclonal antibody to immediately crosslink any biosynthetic NLGN1 reaching the PM (**Fig. 5A**). In control neurons, we also released NLGN1 for 2h, but NLGN1 surface labeling was performed after fixation, before permeabilization (**Fig. 5A**). Crosslinking showed that NLGN1 is directly inserted into both dendritic and AIS PM (**Fig. 5B**), reinforcing that distinct vesicle pools sort NLGNs to dendrites versus the AIS. We analyzed the distribution of NLGN1 at the cell surface by measuring the Polarity Index between dendrite and the AIS. We observed an enrichment of NLGN1 at the AIS PM upon crosslinking, compared to control neurons (**Fig. 5B, D**). This suggests that NLGN1 is retrieved from the AIS by endocytosis or lateral diffusion. We applied the same assay to RUSH-EGFP-NLGN2, which also showed its insertion into dendritic and AIS PM (**Fig. 5C**). However, the crosslinking did not strongly affect the presence of NLGN2 at the AIS PM compared to control conditions (**Fig. 5C, D**). Moreover, NLGN2 was slightly more polarized towards the AIS PM than NLGN1 in control conditions (**Fig. 5C, D**). This indicates that, compared to NLGN1, the presence of NLGN2 at the AIS PM is more stable indicated by a smaller polarity shift upon crosslinking (**Fig. 5E**). As a control, we applied the crosslinking method to biosynthetic TFR, which was inserted at dendritic but not AIS PM (**Fig. S5A**).

**Figure 5.**
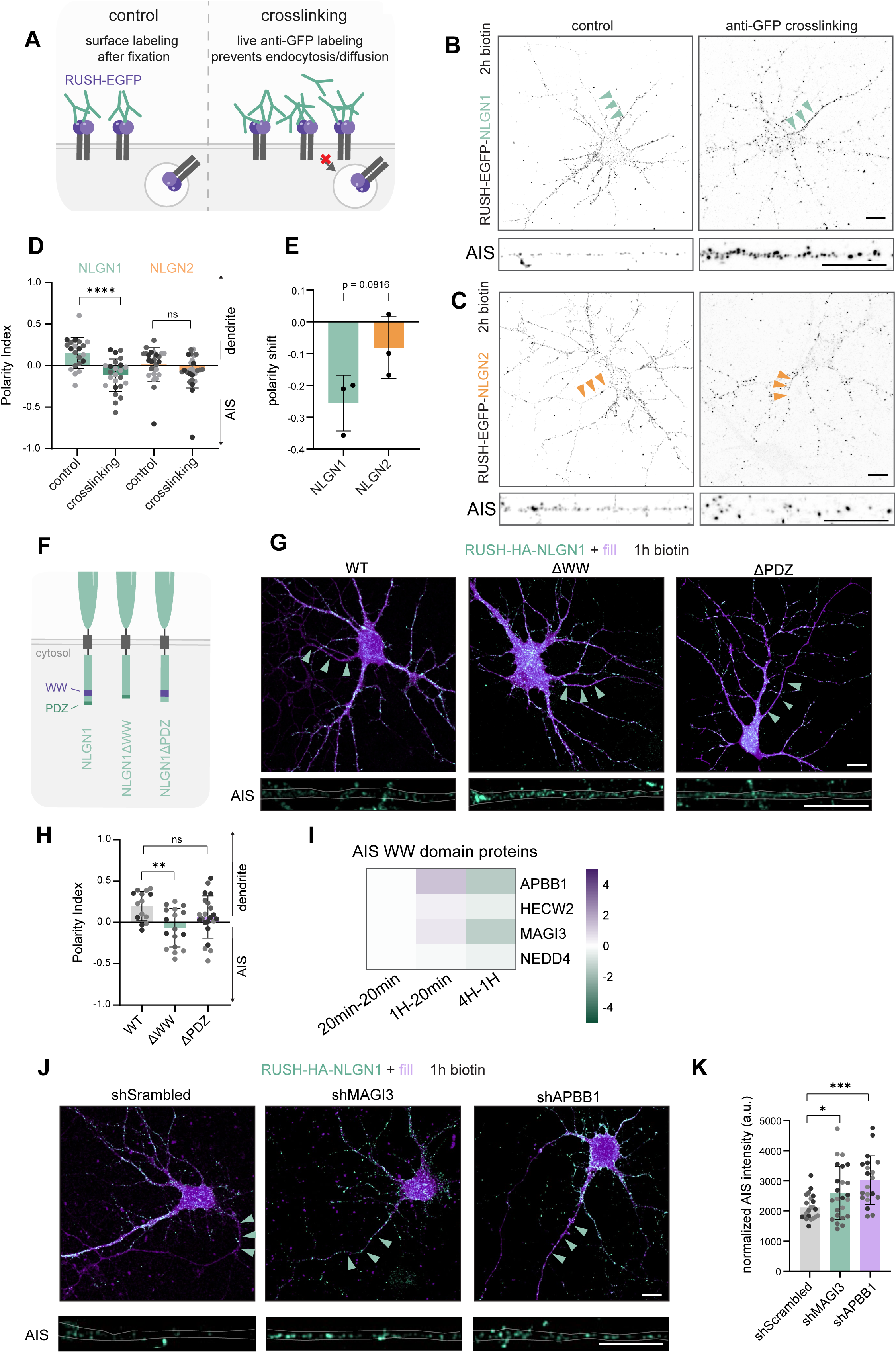
NLGN1 and NLGN2 are inserted into the AIS PM transiently and stably, respectively, with NLGN1 removal mediated by endocytosis, APBB1, MAGI3. (A) Schematic showing that live crosslinking with polyclonal anti-GFP antibodies prevents lateral diffusion or endocytosis of cargo inserted at the PM. (B-E) Representative overview images and AIS crops of DIV15 neurons expressing RUSH-EGFP-NLGN1 (B) or RUSH-EGFP-HA-NLGN2 (C), live incubated with biotin for 2h in the presence of anti-GFP antibody (crosslinking) or stained for GFP after fixation (control). Quantification of the Polarity Index (D). Quantification of the shift in polarity between control and crosslinking per experiment (E). (F) Schematic depicting WW- and PDZ-domain deletions in the C-terminus of NLGN1. (G-H) Representative images of neurons expressing RUSH-HA-NLGN1 WT, ΔWW or ΔPDZ, live incubated with biotin for 1h and surface labeled for HA (G). Quantification of Polarity Index (H). (I) Heat map of WW domain proteins present at the AIS, in which the log2 Fold Change across time points is plotted. (J-K) Representative images of neurons expressing RUSH-HA-NLGN1, tdTomato (fill) and shScrambled, shMAGI3 or shAPBB1, live incubated with biotin for 1h and surface labeled for HA (J). Quantification of AIS intensity, normalized to the intensity in the soma (K). Data points represent individual neurons (except in E), color indicates individual experiment. Kruskal-Wallis test with Dunn’s correction (D), unpaired t test (E), one-way ANOVA with Dunnett’s correction (H), Brown-Forsythe and Welch ANOVA test with Dunnett’s correction (K). ns: no significant, *p < 0.05, **p < 0.01, ***p < 0.001, ****p < 0.0001. Green arrows indicate the AIS. Scale bars represent 10 µm. See also Fig. S5.

Next, we wanted to determine the mechanism involved in NLGN1 removal from the AIS PM. Previously, it was shown that endocytosis at the AIS maintains neuronal polarity by preventing diffusion of somatodendritic cargoes from the somatic to the axonal PM^35^. Since we observed the direct but transient targeting of biosynthetic NLGNs to the AIS PM, we wondered whether endocytosis mediates its retrieval. For this, we released RUSH-HA-NLGN1 from the ER for 1h, while blocking endocytosis with Dynasore and labeling the surface pool. Dynasore treatment increased the amount of RUSH-HA-NLGN1 present at the AIS PM after 1 hour release (**Fig. S5B, C**). This indicates that endocytosis plays a role in the removal of NLGN1 from the AIS PM.

We next set out to investigate which proteins mediate the retrieval of NLGN1. To narrow down the search, we first assessed which domains within NLGN’s intracellular domain are involved. We made NLGN1 mutant constructs lacking either the WW-binding domain or the PDZ ligand (**Fig. 5F**), which are known to be involved in regulating the surface expression of NLGN1 in dendrites^36,37^. A WW-binding domain is typically a proline-rich region recognized by a WW-domain, which has two conserved tryptophans. A PDZ-ligand is a short C-terminal peptide recognized by a PDZ-domain. Removal of the WW-binding domain significantly increased surface distribution of NLGN1 at the AIS, whereas removal of the PDZ ligand led to a small but not significant increase (**Fig. 5G, H**). Further analysis of our MS dataset revealed 11 proteins containing WW-domains at the 1h time point. Upon cross-referencing to an available AIS proteome dataset^32^, we found that of those 11 WW-domain proteins, APBB1, HECWecw2, MAGI3 and NEDD4 are present at the AIS (**Fig. 5I**). APBB1 is a cargo adaptor protein for amyloid precursor protein (APP). It mediates its endocytosis via LRP1 (also present in our proteomic data)^38^. MAGI3 is a scaffolding protein that can interact with NLGN1 via both its WW- and PDZ-domains. MAGI3 has been implicated in the internalization of 5-HT2AR, suggesting a similar role in the internalization of NLGN1^39^. We found that knockdown of APBB1 and MAGI3 increased biosynthetic NLGN1 localization at the AIS PM, indicating their involvement in the endocytosis of NLGN1 (**Fig. 5J, K; Fig. S5D-F**).

Together, these results show that biosynthetic NLGN1 and NLGN2 are transiently or stably inserted into the AIS PM, respectively. NLGN1 is removed from the AIS PM by APBB1- and MAGI3-mediated endocytosis.

### NLGN2, but not NLGN1, predominantly localizes to inhibitory synapses at the AIS

Because of their established role in dendritic synapses, it is possible that these NLGNs at the AIS are integrated into axo-axonic synapses. These inhibitory synapses play a crucial role in homeostatic plasticity^40^. To investigate the presence of endogenous NLGN1 and NLGN2 at these inhibitory synapses, we generated NLGN1 (HiUGE^43^) and NLGN2 (ORANGE^44,45^) knock-ins. In mature neurons (DIV21), the dendritic distribution of both NLGN knock-ins is as reported; NLGN1 colocalizes with the excitatory postsynaptic marker Homer1, while NLGN2 colocalizes with the inhibitory postsynaptic marker gephyrin (GEPH) (**Fig. S6A, B**). Interestingly, we observed the AIS decorated by both NLGN1 and NLGN2 puncta at the steady state (**Fig. 6A-C**). Consistent with our results of more stable biosynthetic NLGN2 than NLGN1 at the AIS PM (**Fig. 5A-E**), we measured a higher number of NLGN2 puncta than NLGN1 at the AIS (**Fig. 6A-C**). Similar distribution of NLGN1 puncta at the AIS was observed in a NLGN1 ORANGE knock-in tagged at the C-terminus (**Fig. S6C**). NLGN2 showed very strong colocalization with both the inhibitory presynaptic marker VGAT (**Fig. 6A-D**) and the inhibitory postsynaptic marker GEPH (**Fig. S6B**). This strongly suggests that, similar to dendritic synapses, NLGN2 localizes to inhibitory synapses at the AIS. We observed some colocalization of NLGN1 and the excitatory postsynaptic marker Homer1 at the very proximal AIS; however, NLGN1 puncta in the rest of the AIS were devoid of Homer1 signal (**Fig. S6A**).

**Figure 6.**
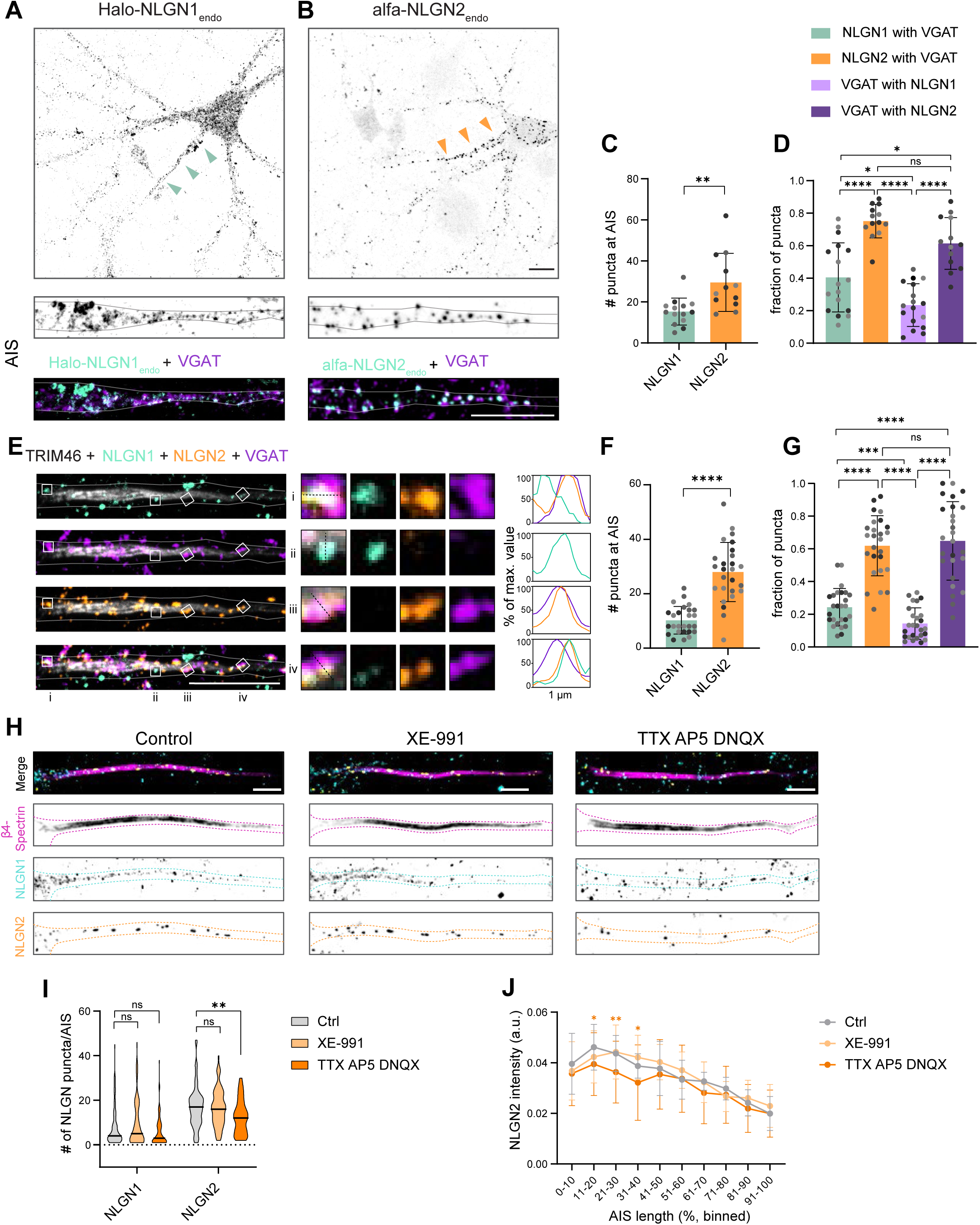
NLGN2, but not NLGN1, predominantly occupies inhibitory synapses at the AIS and is regulated by network activity. (A-D) Representative images of soma and AIS of fully mature neurons (DIV21) with NLGN1 endogenously tagged with HaloTag (A) and NLGN2 endogenously tagged with 3xalfa (B) at the N-terminus and stained for VGAT. Quantification of the number of NLGN puncta at the AIS (C). Quantification of the fraction of NLGN1, NLGN2 and VGAT puncta colocalizing with VGAT, NLGN1, and NLGN2 (D). (E-G) Representative image of the AIS of neurons stained for TRIM46, NLGN1, NLGN2 and VGAT. Line graphs show the intensity as percentage of the maximum value (E). Quantification of the number of NLGN puncta at the AIS (F). Quantification of the fraction of NLGN1, NLGN2 and VGAT puncta colocalizing with VGAT, NLGN1 and NLGN1 (G). (H-I) Representative image of the AIS of neurons stained for β4-spectrin, NLGN1 and NLGN2, treated with either XE-991, a Kv7 channel (KCNQ) blocker, or tetrodotoxin (TTX), AP5, and DNQX for 48h. Scale bars represent 5 µm (H). Quantification of the number of NLGN puncta at the AIS (I). Intensity distribution of NLGN2 along the AIS (J). Data points represent individual neurons; colors indicate individual experiments. Mann-Whitney test (C), Brown-Forsythe and Welch ANOVA with Games-Howell’s correction (D), paired t test (F), RM one-way ANOVA with Tukey’s correction (G). two-way ANOVA with Dunnett’s correction, N=3 independent experiments, n=61-72 cells (I). two-way ANOVA with Tukey’s correction. N=3 independent experiments, n = 30 cells (J). Arrows indicate the AIS. Scale bars represent 10 µm unless otherwise indicated. See also Fig. S6.

Next, we aimed to directly compare NLGN1 and NLGN2 distribution within the same neuron. Thus, we performed simultaneous NLGN labeling with antibodies. Using the immunolabeling approach, we confirmed the greater abundance of NLGN2 puncta compared to NLGN1 at the AIS (**Fig. 6E, F**). We found that ∼ 30% of NLGN1 puncta colocalized with NLGN2 (**Fig. 6E; Fig. S6D, E**), and ∼ 25% colocalized with VGAT (**Fig. 6E, G**). On the other hand, we found that ∼ 15% of NLGN2 puncta colocalized with NLGN1 (**Fig. 6E, S6E**), while its colocalization with VGAT was much more pronounced (∼ 60%; **Fig. 6E, G**).

The AIS has been shown to undergo homeostatic plasticity in response to changes in neuronal activity^46–48^, with alterations in axo-axonic synapses number^40^. Since NLGN2 is directly and stably targeted to inhibitory axo-axonic synapses at the AIS, we wondered whether this would be affected by changes in neuronal network activity. Therefore, we compared the distribution of both NLGN1 and NLGN2 at the AIS in the context of different network activity states. We chronically increased excitability with XE-991 to block potassium channel Kv7^49^ or silenced neuronal networks using tetrodotoxin (TTX), AP5 and DNQX^50–54^ to block voltage-gated sodium channels, NMDARs, and AMPAR for 48h. The XE-991 treatment increases excitability by blocking the Kv7-mediated outward current, while TTX and AP5/DNQX produce a silent network state by blocking sodium channel opening and preventing glutamate receptor binding, respectively. The application of the treatment did not cause AIS structural plasticity, as measured by β4-Spectrin length, nor distance from the soma (**Fig. 6H; Fig. S6F, G**). Interestingly, we found that the number of NLGN2, but not NLGN1 puncta was significantly decreased after the network silencing treatment (**Fig. 6H, I**), suggesting that NLGN2 AIS distribution could be dynamically regulated by activity. Inhibitory synapse positioning along the AIS length has been shown to be important for strength of inhibition^55^. To determine how network activity remodels NLGN2 distribution within the AIS, we plotted NLGN2 intensity and observed that the proximal AIS enrichment of NLGN2 puncta is significantly reduced by network silencing (**Fig. 6J**).

In conclusion, we show the AIS delivery of both NLGN1 and NLGN2 in mature neurons. NLGN2, but not NLGN1, predominantly localized to axo-axonic synapses in an activity-dependent manner, suggesting a role in axo-axonic synaptic plasticity.

### NLGN puncta localize to the AIS in cortical organotypic slice cultures

While primary neuron cultures are ideal for manipulating and visualizing subcellular processes such as biosynthetic trafficking, their circuitry lacks the organization and complexity of native tissue. We therefore sought to characterize the distribution of NLGNs at the AIS of mouse cortical organotypic slice cultures (OSCs), which retain a laminar cortical layering and the major neuronal and glial cell types^56^. Endogenous NLGN1 was detected at the AIS of pyramidal neurons in both the upper (2/3) and lower (5) cortical layers. We found that the vast majority of AIS contained NLGN1 puncta; 75% in L2/3 and 78% in L5, localized to the edge of the AIS membrane (**Fig. 7A-D**). While pyramidal neurons in L2/3 and L5 did not differ in number of NLGN1 puncta, the AIS of L5 neurons were longer, which we found to be independent of NLGN1 content (**Fig. 7E-G**) To underscore this, we further found no correlation between AIS length and number of NLGN1 puncta in these neurons (**Fig. S7A**).

**Figure 7.**
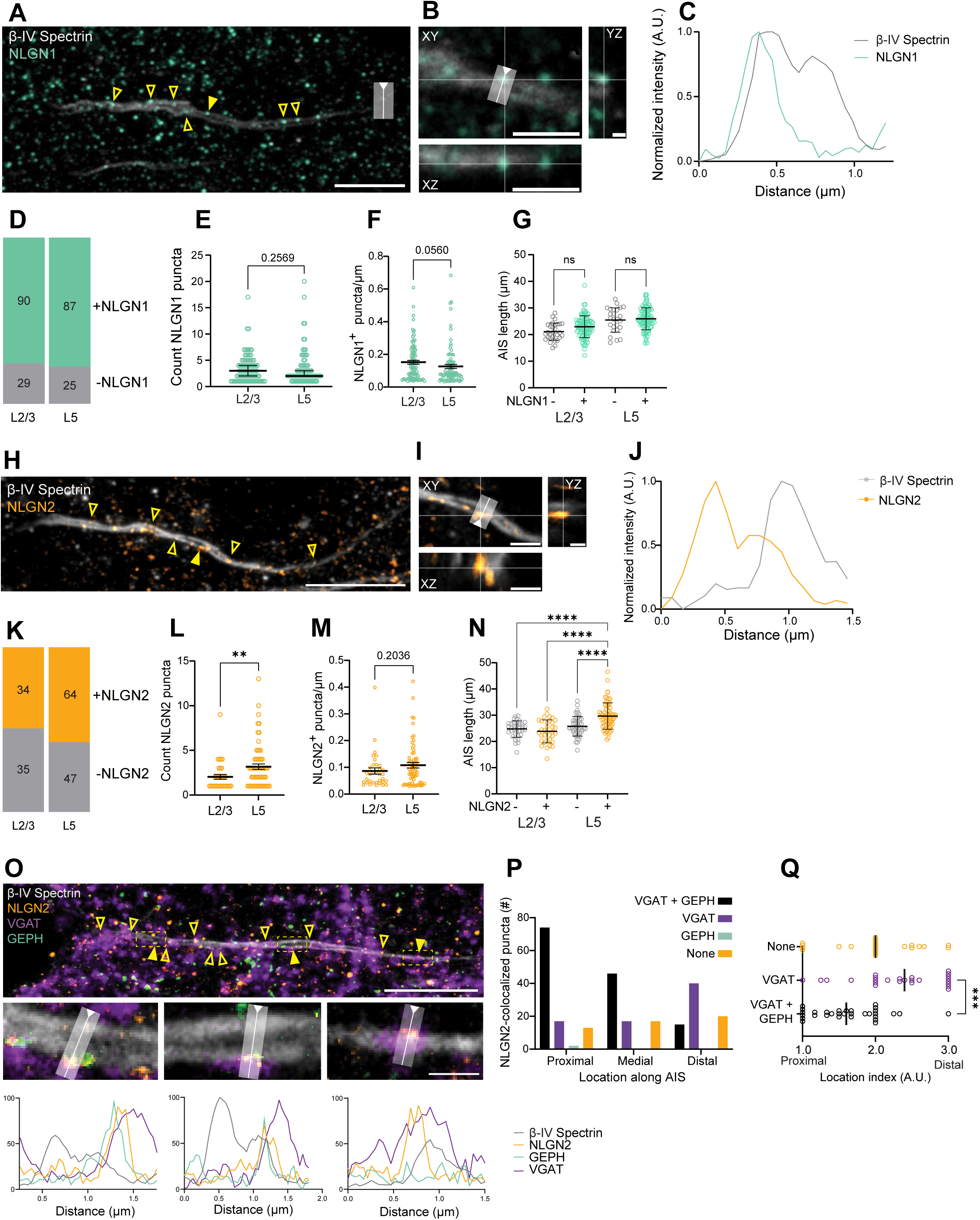
NLGN puncta localize to the AIS in cortical organotypic slice cultures. (A-C) Representative image of immunostained L2/3 pyramidal neuron AIS from DIV19 OSC; (B). Scale bar 10 µm (A). Crop and orthogonal projections of A, showing colocalization of NLGN1 with βIV-Spectrin in three dimensions. Fixation crosses indicate punctum location, box and arrowed line show line profile acquired in C. (B). Line profile (C) (D-G) Fraction of AIS containing NLGN1 compared to those without, N = 4 slices (D). Count (E) and density (F) of NLGN1 puncta in AIS of L2/3 versus L5. Length of AIS with and without NLGN1 in L2/3 and L5 (G). (H-J) Representative image of immunostained L5 pyramidal neuron AIS from DIV19 OSC. Scale bar 10 µm (H) Crop and orthogonal projection of H, fixation crosses indicate punctum location; box and arrowed line indicate line profile in J (I). Line profile (J). (K-N) Fraction of AIS containing NLGN2 compared to those without, N = 4 slices (K). Count (L) and density (M) of NLGN2 puncta in AIS of L2/3 versus L5. Length of AIS with and without NLGN2 in L2/3 and L5; (N). (O-Q) Representative image of immunostained L5 pyramidal neuron from DIV19 OSC. Top, complete AIS; boxes indicating areas cropped out below. Box and arrowed line show line profiles below (O). Differential distribution and composition of NLGN2-colocalized puncta along the AIS (P); Location index (arbitrary units) of solitary NLGN2+ puncta (None), NLGN2+/VGAT+ puncta and NLGN2+/VGAT+/GEPH+ puncta per AIS (Q). N = 2 slices. Yellow triangles indicating colocalized puncta, with solid triangle pointing to respective crops. Scale bars of overview images represent 10 µm, scale bars in zooms represent 2 µm, or 1 µm (YZ). Mann-Whitney U test (E, L), lognormal Welch’s t test (F, M), Kruskal-Wallis test with Dunn’s correction (G). one-way ANOVA with Tukey’s correction (N). **** P < 0.0001 (R). Mixed-effects analysis with Tukey’s correction. ns: no significant, *p < 0.05, **p < 0.01, ***p < 0.001, ****p < 0.0001.

NLGN2 was also detected at the AIS of upper and lower cortical pyramidal neurons (**Fig. 7H-J**). We likewise found NLGN2 to be localized to the AIS membrane (**Fig. 7J**), but in L5, a larger fraction of the AIS contained NLGN2 than in L2/3; 49% in L2/3 and 58% in L5 (**Fig. 7K**). Moreover, the NLGN2-containing AIS in L5 harbored more NLGN2 puncta than in L2/3 (**Fig. 7L**), but this was not significant upon correction for the difference in AIS length (**Fig. 7M**). Nevertheless, we found that NLGN2-containing AIS were longer than AIS lacking NLGN2 (**Fig. 7N**), and there was a positive correlation between the number of NLGN2 puncta and AIS length in layer 5 (**Fig. S7B**).

Lastly, we investigated the distribution of NLGN2 at axo-axonic synapses. Therefore, we labeled OSCs for NLGN2, VGAT and GEPH. Just as in cultured dissociated neurons, we found NLGN2 present at functional synapses, labeled by both pre- and postsynaptic markers (**Fig. 7O**). Interestingly, we found that NLGN2^+^ puncta in functional synapses were predominantly localized at the proximal AIS, whereas the more distal AIS harbored more VGAT+/NLGN2+/GEPH-puncta (**Fig. 7P-Q**). The number of orphan NLGN2 puncta (without VGAT or GEPH) was evenly distributed along the AIS (**Fig. 7P-Q**).

Taken together, we show that NLGN1 and NLGN2 localize to the AIS of pyramidal neurons in cortical organotypic slice cultures. The distribution of NLGN2 at axo-axonic synapses reinforce the idea of a possible role for NLGN2 in tuning the strength and/or number of axo-axonic synapses for homeostatic plasticity.

### NLGN stability at the AIS is determined by the ectodomain

Our results demonstrate that NLGN2, but not NLGN1, is a functional component of axo-axonic synapses in primary neurons and OSCs; nevertheless, we consistently observe the targeting and transient localization of NLGN1 at the AIS. The ectodomain of NLGNs determines their ability to enhance excitatory/inhibitory synaptic function^16,57^. However, it is not clear how the intra- vs. ecto-domains are involved in NLGN distribution and retention at the AIS PM. To study this, we made RUSH chimera constructs of the extracellular domain of NLGN1 with the intracellular domain of NLGN2, and vice versa (**Fig. 8A**). We transfected these RUSH-chimera constructs and performed the live antibody crosslinking assay, while releasing the cargo with biotin for 2h. We observed that the NLGN1-NLGN2 chimera, bearing the extracellular domain of NLGN1, behaved similarly to wild-type NLGN1; specifically, its localization at the AIS was enhanced upon crosslinking, indicating its transient localization at the PM (**Fig. 5A-E**; **Fig. 8B, D, E**). On the other hand, the NLGN2-NLGN1 chimera, containing the intracellular domain of NLGN1, showed limited effect of crosslinking, like wild-type NLGN2 (**Fig. 5A-E**; **Fig. 8C, D, E**). In the same experiment, we also labeled the inhibitory post-synaptic marker GEPH (**Fig. 8F, G**). We found that the NLGN2-NLGN1 chimera colocalizes more with GEPH than the NLGN1-NLGN2 chimera, albeit less strong than with the wild-type NLGN2 (**Fig. 8F, G**). This difference is probably due to the collybistin-binding domain, which provides an additional link to GEPH and is missing from the NLGN1 intracellular domain^58^.

**Figure 8.**
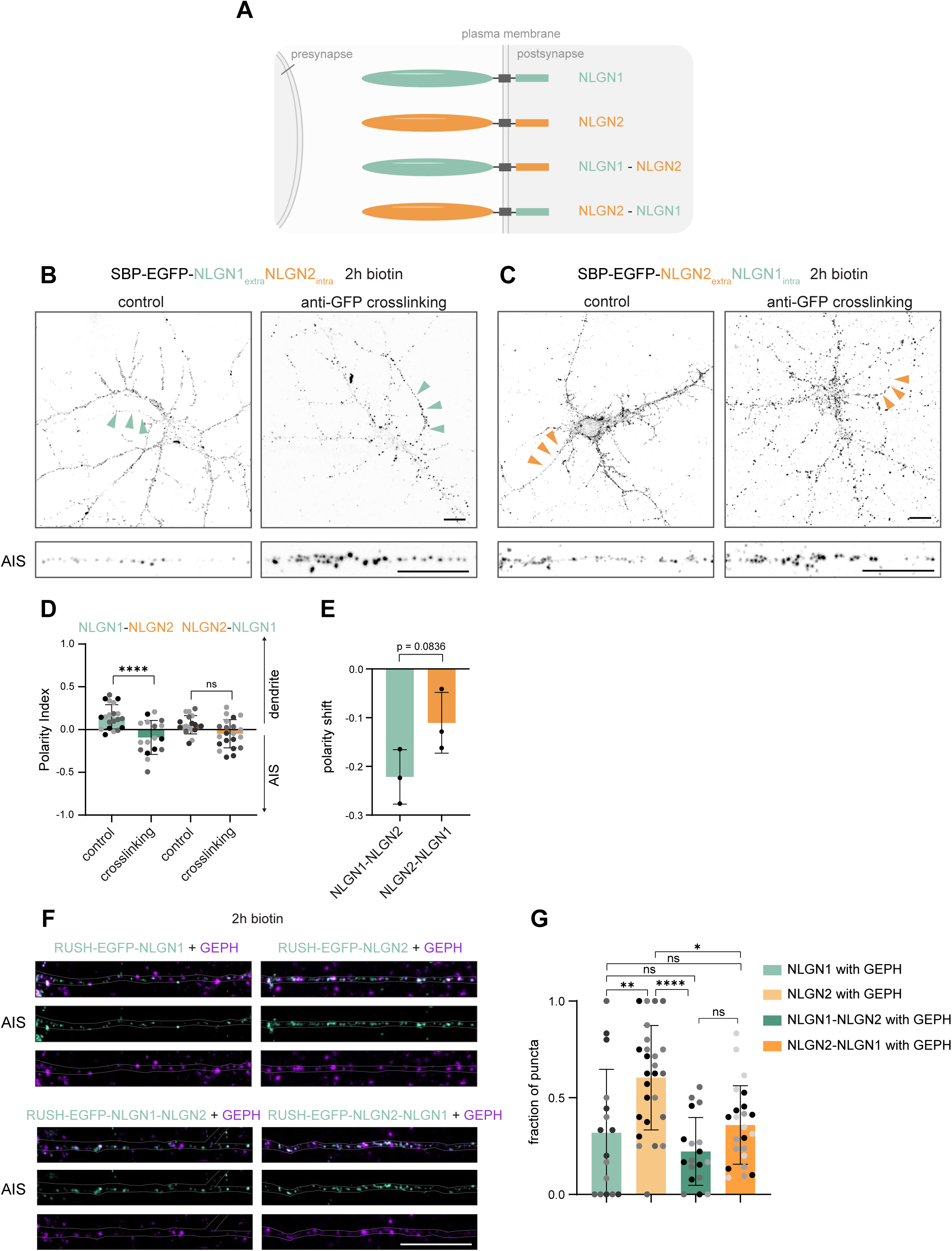
NLGN stability at the AIS is determined by the ectodomain. (A) Schematic depicting wild-type NLGN1 and NLGN2, and chimera with the intracellular domain swapped. (B-C) Representative overview images and AIS crops of DIV15 neurons expressing RUSH-EGFP-NLGN1_extra_NLGN2_intra_ (B) or RUSH-EGFP-HA-NLGN2_extra_-NLGN1_intra_ (C), live incubated with biotin for 2h in the presence of anti-GFP antibody (crosslinking) or stained for GFP after fixation (control). (D-E) Quantification of the Polarity Index (D) Quantification of the shift in polarity between control and crosslinking per experiment (E). (F-G) Representative overview images and AIS crops of neurons transfected with RUSH-EGFP-NLGN1, RUSH-EGFP-NLGN2, RUSH-EGFP-NLGN1_extra_NLGN2_intra_ or RUSH-EGFP-HA-NLGN2_extra_-NLGN1_intra_, incubated with biotin for 2h and stained after fixation, before permeabilization for EGFP, and after permeabilization for GEPH (as in Fig 5B-C and Fig. 8B-C control) (F). Fraction of NLGN puncta colocalizing with GEPH (G). Data points represent individual neurons, colors indicate the individual experiments. Ordinary one-way ANOVA with Sidak’s correction (D), unpaired t test (E), Brown-Forsythe and Welch ANOVA with Holm-Sidak’s correction (G). Scale bars represent 10 µm.

These results indicate that the NLGN extracellular domain determines whether these proteins are retrieved from or stabilized at axo-axonic synapses at the AIS PM.

## Discussion

Neuronal development and function rely on precise regulation of membrane protein sorting to distinct neuronal domains, yet the underlying mechanisms remain incompletely understood^6^. Here, we investigated the trafficking of highly homologous NLGNs, which have distinct localizations and functions at dendritic excitatory and inhibitory synapses. Applying the POTATOMap tool^18^ we dissected the itinerary of biosynthetic NLGN1 identifying co-cargoes and sorting routes. Using genetic engineering and microscopy, we further characterized dendritic vesicle populations and their sorting mechanisms. Interestingly, we elucidated the trafficking and localization of NLGN1 and NLGN2 at the AIS. Because of their highly similar and interchangeable intracellular domains, we propose a model where biosynthetic NLGNs are sorted together to the AIS PM. Only after their insertion at the PM, NLGNs bind pre-synaptic neurexins, thereby ‘detecting’ synapse type. Since the AIS contains only inhibitory synapses, NLGN2 is retained, while NLGN1 is retrieved. These findings open new avenues to explore the role of AIS-localized NLGNs in shaping AIS structural and functional plasticity.

The mechanisms involved in biosynthetic NLGN trafficking were unknown. Here, using POTATOMap, we found a group of PM proteins as potential co-cargoes. Some of these are sorted into distinct vesicles at the TGN by either AP-1 (TFR)^2,3^, or AP-4 (AMPAR)^1^. Although NLGNs lack canonical AP binding motifs, multiple APs appeared in our POTATOMap interactome. It remains unknown whether NLGNs bind these APs in a non-canonical manner at the TGN. By developing a biosynthetic axonal relocation assay for NLGN1, we confirmed co-sorting of biosynthetic NLGN1 with selected co-cargoes. Furthermore, we found NLGN1 distributed across different vesicle types: while TFR and NLGN1 mostly trafficked together, we found a NLGN population that traffics without TFR to the AIS. Possibly, NLGN1+/TFR+ vesicles are sorted by AP-1, while NLGN1+/TFR-vesicles targeting dendrites and the AIS, are sorted by AP-4 and AP-3, respectively. Lastly, we also identified NLGN2 and NLGN3 as potential co-cargoes of NLGN1; however, these were not tested by our axonal relocalization tool due to their potential heterodimerization^59^. Nevertheless, their interchangeable intracellular domains support their co-trafficking.

We further exploited our MS data to investigate mechanisms involved in the sorting of NLGN1 to dendrites. We studied AFTPH and NSG2, two potential players in regulating protein sorting at the TGN^21–23^. We found that AFTPH and NSG2 were both required for polarized trafficking of NLGN1 to dendrites. Their knockdown caused missorting of NLGN1 to the axon, resembling the effect of AP-1 disruption on TFR mislocalization^22^. AFTPH is known to form a complex with HEATR5B and SYNRG to facilitate AP-1 function^60^. We also observed these two proteins in our MS data, suggesting the role of this complex in regulating AP-1-mediated polarized trafficking to dendrites. On the other hand, NSG2 has been poorly studied; by homology with NSG3, it is predicted to interact with clathrin^24,61^. Although it distributes to the TGN^23,62^, and its knockdown impairs polarized trafficking of biosynthetic NLGN1, its exact function within the biosynthetic pathway remains unclear. It would be important to examine whether NSG2 facilitates polarized sorting to dendrites via AP-1. Its interaction with AP-4 is unlikely as AP-4 is considered a non-clathrin-associated adaptor^5^. Lastly, we found that depletion of the kinesin-3 KIF1A reduced trafficking of NLGN1 into dendrites. KIF1A transports dense core vesicles (DCVs) into dendrites, then liprin-α (PPFIA2) and TANC2, which were also present in our MS data, recruit KIF1A-DCV into spines^26^. It is important to note that dendritic trafficking of NLGN1 was not completely abolished upon KIF1A depletion, thus we do not rule out a role for KIF1B, KIF3A/B or dynein, present in our MS data, for NLGN1-positive vesicle transport to dendrites.

In addition to the sorting of NLGNs to dendrites, we found that they are also sorted to the AIS PM. A previous study showed the transient targeting of NLGN1 to the AIS PM^17^, which we also observed in our MS data, live cell imaging and surface crosslinking experiments for RUSH-NLGN1. For a long time it was believed that exocytosis at the AIS was limited, due to its dense actin-spectrin network, with exocytosis of AIS proteins taking place at the somatic or axonal PM, followed by selective retention at the AIS^63,64^. Recent evidence and our work now challenge this view, showing that exocytosis of cargoes occurs at the AIS^17,65^. We found that the sorting of biosynthetic NLGN-positive vesicles to the AIS is mediated by the axonal motor kinesin-1, yet we rarely observe NLGNs actually moving towards the axon. It is not known how vesicles containing NLGN, or any other AIS-specific protein for that matter, are halted at the AIS and directed to the AIS PM for fusion. Interestingly, the AIS scaffold protein AnkyrinG has been shown to bind kinesin-1^66^, suggesting a possible mechanism for recruiting kinesin-1-mediated AIS-containing vesicles to the AIS for their PM fusion. We determined that NLGN1 is more transiently associated with the AIS PM than NLGN2. Moreover, we showed that NLGN1 is retrieved from the AIS through endocytosis, in a mechanism that depends on its WW binding domain, and to a much lesser extent on its PDZ ligand. By mining our MS dataset for proteins that could bind the WW-binding domain, and which are present at the AIS^32^, we were able to identify two proteins possibly implicated in this process, MAGI3 and APBB1. MAGI3 is interesting as it contains multiple WW and PDZ domains, thereby acting as a signaling platform that senses the local environment and can recruit relevant interactors^39^. APBB1 is a cargo adaptor protein for APP; it mediates its endocytosis via LRP1 (also present in our MS data)^38^. As it appears to affect NLGN1 similarly, it suggests a common pathway for adhesion molecules.

By employing NLGN1 and NLGN2 chimera constructs, we found that NLGN2 stability at the AIS PM depends on its extracellular domain in hippocampal cultures. This domain plays a key role in determining NLGN distribution across inhibitory versus excitatory synapses. As the AIS harbors only inhibitory synapses, NLGN2 is the most dominant paralog here. We can speculate that NLGN1 at the AIS PM encounters less presynaptic binding partners, it is not stabilized and therefore removed. Although we cannot exclude a function for NLGN1 at the AIS, it appears that is mainly trafficked to this compartment because of its similarity to NLGN2. This might seem ineffective, however, in the dendritic context it makes sense to insert all NLGN paralogs non-preferentially at the PM, as their role in synapse assembly means they need to sense the extracellular presynaptic partners, at both excitatory and inhibitory, to then recruit the correct postsynaptic scaffolding and receptors. Nevertheless, the chimeras do not behave entirely like WT NLGN1 and NLGN2, respectively. Therefore, we cannot rule out specific functions for the intracellular domain of each paralog. Lastly, previous research suggested a prominent role for the adhesion molecules Neurofascin, L1CAM and Contactin-1 in organizing axo-axonic synapses and recruiting GEPH at the AIS^41,42,67,68^. No such role has been described for NLGNs before; thus, it would be important to determine the role of NLGNs in GEPH recruitment at the AIS.

Consistent with the more stable recruitment of NLGN2 than NLGN1 to axo-axonic synapses, we found that in the absence of network activity the number of NLGN2 puncta along the AIS reduced, which could hint at a role for NLGN2 in mediating activity-dependent scaling of the inhibitory synapses at the AIS. While increasing activity by blocking Kv7 channels did not lead to an opposite change, future studies would require electrophysiological recordings to more directly determine the contribution of action potential generation to the distribution of NLGN2 at the AIS. Interestingly, we found that while NLGN2 was part of the pre- and postsynaptic complex at the proximal AIS, in the distal AIS it appeared more often with VGAT but without GEPH. It remains to be investigated with live imaging whether these puncta represent remnants of pruned synapses, or perhaps provide a starting point for the formation of new axo-axonic synapses.

In the brain axo-axonic synapses are established by chandelier interneurons which uniquely target the AIS with inhibitory terminals^69^. Although the role of chandelier synapses to the excitability of the AIS, and its action potential output, is debated, recent studies showed that anatomical remodeling of axo-axonic inhibition is involved in learning-induced structural changes of the AIS^70–72^. The present finding that NLGN2 is a part of the axo-axonic synaptic complex may help to elucidate molecular insight into activity-dependent remodeling and its role in AIS plasticity will be an exciting avenue for further investigation. In addition, changes in axo-axonic synaptic innervation patterns have also been linked to epilepsy^73^, but also schizophrenia^74^ and Autism Spectrum Disorder (ASD)^75^, reviewed in ^76,77^. As NLGN2 has also been linked to epilepsy^12,78^ and ASD^79,80^, future studies in neurological and neuropsychiatric disease models may shed light on the NLGN2 function at axo-axonic synapses.

## Acknowledgments

This work was supported by the European Research Council (ERC-StG 950617) to GGF, (ERC-StG 101117227) to AF, by an NWO (Dutch Research Council) VIDI grant (Grant No. VI.Vidi.233,056) to AF, and by funding from the Institute of Chemical Immunology (project ICI000032) to MHPK. We thank Dr. Wouter Droogers and the MacGillavry lab for their help with creating knock-in constructs.

## Author contributions

N.K.: Conceptualization, investigation, formal analysis, methodology, visualization, supervision, writing.

C.H.L.: Investigation, formal analysis, methodology.

V.N., L.H.: Investigation, formal analysis, visualization, writing.

V.V., E.M., P.U., G.L.: Investigation, formal analysis.

M.A.: Supervision, resources.

A.F., M.K.: Funding acquisition, supervision, writing.

G.G.F.: Conceptualization, funding acquisition, supervision, writing.

## Declaration of generative AI and AI-assisted technologies in the writing process

During the preparation of this manuscript, the authors used Perplexity and ChatGPT to improve language clarity. The authors reviewed and edited the content as needed and take full responsibility for the content of the published article.

## Methods

### Key resources table

| REAGENT or RESOURCE | SOURCE | IDENTIFIER |
| --- | --- | --- |
| Rabbit-anti-TRIM46 | van Beuningen et al.<sup>81</sup> | N/A |
| Mouse anti-V5 | Thermo Fisher Scientific | R960-25; RRID:AB_2556564 |
| Rat anti-HA | Roche | 11867423001; RRID:AB_390918 |
| Mouse anti-Pan-Neurofascin external | UC Davis / NIH NeuroMab | 75-172; RRID:AB_2282826 |
| Rabbit anti-GM130 | Abcam | RRID:AB_880266 |
| rabbit anti-ALFA (rHcAb) | NanoTag Biotechnologies | N1583; RRID AB_3075998 |
| Guinea Pig anti-PSD95 | Kim et al<sup>82</sup> | N/A |
| Mouse anti-GABAA | Millipore | MAB341; RRID:AB_11214320 |
| Rabbit anti-Halo | Promega | G9281; RRID:AB_713650 |
| Mouse anti-GM130 | BD Biosciences | 610823; RRID:AB_398142 |
| chicken anti-GFP | Aves Labs | GFP-1020; RRID AB_2307313 |
| chicken anti-VGAT | Synaptic Systems | 131009; RRID:AB_2943610 |
| mouse anti-gephyrin | Synaptic Systems | 147011; RRID:AB_887717 |
| mouse anti-AnkyrinG | Thermo Fisher Scientific | 33-8800; RRID:AB_2533145 |
| Rabbit anti -βIV-Spectrin | Homemade | Epitope: aa 2237-2256 of human βIV-Spectrin |
| Mouse anti-NLGN1 | SantaCruz | sc-365110, RRID: AB_10708272 |
| Mouse anti-NLGN2 | Synaptic Systems | Cat#129511, RRID: AB_2619813 |
| Guinea pig anti-ankyrinG | Synaptic Systems | Cat# 386004, RRID:AB_2725774 |
| Rabbit anti-VGAT | Merck-Millipore | Cat#AB5062P, RRID:AB_2301998 |
| Chicken anti-gephyrin | Synaptic Systems | Cat#147009; RRID:AB_2943527 |
| Goat anti mouse Alexa 405 | Thermo Fisher Scientific | A-31553; RRID:AB_221604 |
| Goat anti mouse Alexa 488 | Thermo Fisher Scientific | A32723, RRID: AB_2633275 |
| goat anti mouse-IgG1 Alexa 647 | Thermo Fisher Scientific | A21240; RRID:AB_2535809 |
| goat anti rabbit Alexa 405 | Thermo Fisher Scientific | A-31556/ RRID:AB_221605 |
| rabbit anti-mouse immunoglobulins/HRP | Agilent | P0260; RRID:AB_2636929 |
| goat anti-mouse IgG2a Alexa 594 | Thermo Fisher Scientific | A-21135, RRID:AB_2535774 |
| goat anti-rabbit AlexaFluor 568 | Thermo Fisher Scientific | A11036, RRID: AB_10563566 |
| goat anti-rabbit AlexaFluor 647 | Thermo Fisher Scientific | A21245, RRID: AB_2535813 |
| goat anti-rat Alexa 488 | Thermo Fisher Scientific | A-11006; RRID:AB_2534074 |
| goat anti-rat Alexa 568 | Thermo Fisher Scientific | A-11077; RRID:AB_2534121 |
| goat anti-rabbit Alexa 488 | Thermo Fisher Scientific | A-11034; RRID:AB_2576217 |
| goat anti-chicken Alexa 405 | Abcam/BioConnect | ab175675; RRID:AB_2810980 |
| goat anti-chicken Alexa 568 | Thermo Fisher Scientific | A11041; RRID:AB_2534098 |
| Goat anti-guinea pig Alexa 568 | Thermo Fisher Scientific | A11075, RRID: AB_141954 |
| goat anti-guinea pig Alexa 647 | Thermo Fisher Scientific | A-21450; RRID AB_2535867 |
| **Chemicals, peptides and recombinant proteins** |  |  |
| Streptavidin, Alexa-Fluor555 | Thermo Fisher Scientific | S21381; RRID:AB_2307336 |
| FluoTag®-X2 anti-ALFA<br>AZDye568 | NanoTag<br>Biotechnologies | N1502 |
| Biotin-phenol | Iris Biotech | LS.3500 |
| Hydrogen Peroxide | Sigma-Aldrich | H1009 |
| Lipofectamine2000 | Thermo Fisher Scientific | 11668019 |
| Polyethylenimine (PEI MAX) | Polysciences | 24765 |
| Fluoromount-G Mounting<br>Medium | Thermo Fisher Scientific | 00-4958-02 |
| Pierce Streptavidin Magnetic<br>beads | Thermo Fisher Scientific | 88816 |
| Trolox | Sigma-Aldrich | 238831 |
| Sodium L-Ascorbate | Sigma-Aldrich | 0000374819 |
| Sodium Azide | Sigma-Aldrich | K43547688 |
| TCEP | Sigma |  |
| IAA | Sigma |  |
| DTT | Sigma |  |
| Bis-Acrylamide 40% | Bio-Rad | #1610144 |
| TEMED | Bio-Rad | 1610800 |
| APS | Sigma-Aldrich | A3678 |
| anhydrous-DMSO | Thermo Fisher Scientific | D12345 |
| Triton-X-100 | Sigma-Aldrich | 93433 |
| 1x protease inhibitors | Roche |  |
| Rapalog (A/C<br>Heterodimerizer) | Takara Bio | 635057 |
| Biotin | Sigma-Aldrich | B4501 |
| Mix-n-Stain CF640R<br>antibody labeling kit | Biotium | 92245 |
| JFX554 | Lavid Lab, Janelia | N/A |
| Dynasore | Tocris | 2897 |
| Tetrodotoxin | SanBio B.V. | SIH-603-1MG |
| DNQX | Tocris | 0189 |
| AP5 | SanBio B.V. | 14539-10 |
| XE-991 | Tocris | 2000/10 |
| **Experimental models: Cell lines** |  |  |
| HEK293T | ATCC | CRL-3216 |
| **Experimental models: organisms/strains** |  |  |
| Rat (Wistar) | Janvier | N/A |
| Mouse (C57BL/6) | In-house | N/A |
| **Recombinant DNA** |  |  |
| FUGW | Kind gift from Dr. David<br>Baltimore | Addgene; plasmid #14883<sup>83</sup> |
| pLKO.1 puro | Kind gift from Dr. Bob<br>Weinberg | Addgene; plasmid #8453<sup>84</sup> |
| psPAX2 | Kind gift from Dr. Didier<br>Trono | Addgene; plasmid #12260 |
| PMD2.G | Kind gift from Dr. Didier Trono | Addgene; plasmid #12259 |
| Halo-TfR-RUSH | Kind gift from Dr. Jennifer Lippincott-Schwartz | Addgene; plasmid #166905<sup>85</sup> |
| PEGFP(A206K)-C1 | Kind gift from Dr. Jennifer Lippincott-Schwartz | N/A |
| pOC1 FLAG-SpCas9 | Kind gift from Dr. Harold MacGillavry | Addgene #183423<sup>45</sup> |
| pCamK Homer1c-mRuby3 | Kind gift from Dr. Harold MacGillavry | N/A |
| ptdTomato-C1 | Kind gift from Dr. Harold MacGillavry | N/A |
| pJet1.2 Halotag donor | Kind gift from Dr. Harold MacGillavry | N/A |
| pcDNA3_HA-KLC-TPR6 | Kind gift from Dr. Kristen Verhey | Verhey et al.<sup>34</sup> |
| TFR-GFP | Kind gift from Dr. Casper Hoogenraad |  |
| HA-GLUR2 | Kind gift from Dr. Casper Hoogenraad |  |
| pNICE-NL2A | Kind gift from Dr. Peter Scheiffele | Addgene; plasmid #15259<sup>86</sup> |
| pFSW_67 | Kind gift from Dr. Harold MacGillavry | N/A |
| Str-KDEL_SBP-EGFP-Ecadherin | Kind gift from Dr. Franck Perez | Addgene plasmid #65286<sup>19</sup> |
| 3xalfa donor vector | Kind gift from Dr. Harold MacGillavry | 3xalfa donor vector |
| pX333 DRS-1-only FLAG-Cas9 | Kind gift from Dr. Harold MacGillavry | pX333 DRS-1-only FLAG-Cas9 |
| pCAMK GFP | Kind gift from Dr. Harold MacGillavry | pCAMK GFP |
| smHA donor vector | Kind gift from Dr. Harold MacGillavry | smHA donor vector |
| pFSW67 | Kind gift from Dr. Harold MacGillavry |  |
| GFP-BICD-N-FRB | Kind gift from Dr. Lukas Kapitein |  |
| mCherry-KIF5A-MD-Strep | Farías et al.<sup>87</sup> |  |
| FUGW_Strep-KDEL_SBP-NLGN1-V5-APEX2 | This study |  |
| FUGW_Strep-KDEL-mNG | This study |  |
| FUGW_SBP-NLGN1-V5-APEX2 | This study |  |
| FUGW_StrepKDEL_mNG-V5-TFR-SBP | This study |  |
| CMV_Strep-KDEL_SBP-NLGN1-Halo-FKBP | This study |  |
| mCherry-KIF5A-FRB | This study |  |
| HA-KIF5A-MD-FRB | This study |  |
| pFSW-StrepKDEL | This study |  |
| pFSW_Strep-KDEL_SBP-NLGN1-mNG | This study |  |
| pFSW_StrepKDEL_SBP-HA-NLGN1 | This study |  |
| pFSW_StrepKDEL_SBP-HA-mNG-NLGN1 | This study |  |
| pFSW_Str-KDEL_SBP-EGFP-HA-NLGN2 | This study |  |
| pFSW_StrepKDEL_SBP-HA-Nlgn1dWW | This study |  |
| pFSW_StrepKDEL_SBP-HA-Nlgn1dPDZ | This study |  |
| pFSW_StrepKDEL_TfR-EGFP-SBP | This study |  |
| pFSW_StrKDEL_SBP-EGFP-HA-NLGN2ec-NLGN1ic | This study |  |
| pFSW_StrKDEL_SBP-EGFP-NLGN1ec-NLGN2ic | This study |  |
| pFSW_StrepKDEL_SBP-EGFP-NLGN1 | This study |  |
| chick-beta-actin_TEN2-EGFP | This study |  |
| pX333_DRS-1-only FLAG-Cas9 NLGN1 | This study |  |
| pAAV_pOC2_NLGN1-ORANGE_smHA | This study |  |
| pAAV_pOC2_NLGN2-ORANGE_3xalfa | This study |  |
| pLKO.1-puro-shScrambled | Li et al.<sup>18</sup> |  |
| pLKO.1-puro-shKIF1A | Farías et al.<sup>89</sup> |  |
| pLKO.1-puro-shNSG2 | This study |  |
| pLKO.1-puro-shAFTPH | This study |  |
| pLKO.1-puro-shMAGI3 | This study |  |
| pLKO.1-puro-shAPBB1 | This study |  |
| **Software and algorithms** |  |  |
| QuantStudio Software V1.3 | Applied Biosystems | https://quantstudio-design-analysis-software.software.informer.com/1.3/ |
| MaxQuant (v2.0.1.0) | MPI of Biochemistry |  |
| Perseus (v1.6.15.0) | MPI of Biochemistry | https://maxquant.net/perseus/ |
| R with R Studio (v4.5.0) | Bell Laboratories | https://www.r-project.org/ |
| Visual Studio Code (v1.109) | Microsoft | https://code.visualstudio.com/ |
| Fiji/ImageJ (v2.16.0/1.54p) | NIH | https://imagej.net/software/imagej/ |
| GraphPad Prism (v10.1.1) | GraphPad | https://www.graphpad.com/ |
| CellProfiler 4.2.8. | Stirling et al.<sup>88</sup> | https://cellprofiler.org/ |

### Resource availability

#### Lead contact

Further information and requests for resources, plasmids and reagents should be directed to and will be fulfilled by the lead contact Ginny G. Farías.

#### Materials availability

Plasmids in this study will be available upon request as of the date of publication.

#### Data and code availability

The mass spectrometry proteomics data generated in this study will be deposited to the ProteomeXchange Consortium via the PRIDE partner repository. The code used for mass spectrometry data analysis in this study is available on the Farías lab GitHub repository.

### Experimental model and study participant details

#### Animals

All animal experiments were approved by the “Centrale Commissie Dierproeven” (Central Commission for Animal Experiments) of the Netherlands Government and were performed according to the Netherlands Law on Animal Research (Wet op Dierproeven) in full agreement with the Directive 2010/63/EU with local approval by and under supervision of the Animal Welfare Body UU, VU, Amsterdam UMC, and Netherlands Institute for Neuroscience. The animal protocol has been evaluated and approved by the national CCD authority–Female pregnant Wistar rats were obtained from Janvier, and embryos (both genders) at embryonic (E)18 stage of development were used for primary cultures of hippocampal and cortical neurons. Mice for organotypic slices were obtained from in-house breeding facilities. The animals, pregnant females and embryos have not been involved in previous procedures.

#### Primary neuron culture and transfection

Hippocampi or cortices from embryonic day 18 rat brains were dissected and dissociated with trypsin for 15 min and plated on coverslips coated with poly-L-Lysine (37.5 µg/ml) and laminin (1.25 µg/ml) at a density of 100,000/well (12-well plates) for imaging; or 1,000,000/well (6-well plates) for proteomics. The day of neuron plating corresponds to day-in-vitro 0 (DIV0). Neurobasal medium (NB) supplemented with 2% B27 (GIBCO), 0.5 mM glutamine (GIBCO), 15.6 µM glutamate (Sigma), and 1% penicillin/streptomycin (GIBCO) was used to maintain neurons incubated under controlled temperature and CO2 (37 °C, 5% CO2). Hippocampal neurons were transfected between DIV7-14 using Lipofectamine 2000 (Invitrogen). For knockdown experiments, they were transfected at DIV4. For knock-in experiments, they were transfected between DIV2-DIV4. Briefly, DNA (10-1000 ng/well) was mixed with 1-3 µL of lipofectamine in 200 µL of NB with 0.5 mM glutamine. DNA/lipofectamine mixture was incubated at room temperature for 20 min. This was added to neurons in ca. 300 µL of their original medium and incubated for 1h at 37 °C, 5% CO2. For knock-in experiments, transfection mix was incubated for 1.5h. For all RUSH experiments, all medium was removed from the neurons and RUSH medium (NB medium containing 2% biotin free B27, 0.5 mM glutamine and 1% penicillin/streptomycin) was added. B27 was subjected to biotin removal using ZebaTM Dye and Biotin Removal Spin Columns (Invitrogen) according to manufacturer’s protocol. In cases when this lead to incomplete biotin removal, we supplemented the medium with Pierce™ Avidin (ThermoFisher). For other experiments performed until DIV10, we transferred the coverslips after transfection to 500 µL of their original medium, supplemented with 500 µL fresh NB medium containing 2% biotin free B27, 0.5 mM glutamine and 1% penicillin/streptomycin. For knock-in experiments, they were transferred to 500 µL of their original medium supplemented with 500 µL BrainPhys (BP) containing 2% NeuroCult SM1 (Stem Cell Technologies) and 1% penicillin/streptomycin (BP-complete). For culturing past DIV14, medium was refreshed weekly by removing 400 µL old medium and supplemented with 500 µL BP-complete.

Experiments were performed within 1-day post-transfection (DIV7-10), or for knockdown experiments 3 days post-transfection (DIV7). For knock-in experiments, experiments were performed either at DIV9 or at DIV21.

#### Cortical organotypic slice preparation

Organotypic slices were created from P4-5 neonatal mouse pups as described previously ^56^. In short, mice were decapitated, brains extracted and placed in ice-cold dissection solution consisting of 98% GBSS ((in mM) 1.5 CaCl2, 0.2 KH2PO4, 0.3 MgSO4, 2.7 NaHCO3, 5.0 KCl, 1.0 MgCl2, 137 NaCl, 0.85 Na2HPO4, 5.6 D-glucose), 1% kynurenic acid (from 0.1 M stock), and 1% glucose (from 2.5 M stock), sterile filtered and adjusted to pH 7.2 as well as 320 mOsm (using sucrose). In a biological safety cabinet, hemispheres were cut and individually sliced to 300 µm thin slices using a McIlwain Tissue Chopper. Slices were then placed onto tissue culture inserts (Merck-Millipore, PICMOGR50) within 6-well plates with 1 ml pre-adjusted culturing medium consisting of 47.75% MEM (Thermo Fisher Scientific # 11575032), 25% HBSS (Thermo Fisher Scientific # 24020133), 25% heat-inactivated horse serum (Thermo Fisher Scientific # 26050088), 2% (2.5 M stock) D-glucose, and 1.25% (1 M stock) HEPES (Sigma-Aldrich H3375), sterile filtered and adjusted to pH 7.2 as well as 320 mOsm. Medium was fully replaced three times per week, and once matured to the desired age, slices were fixated using chilled 4% para-formaldehyde (Thermo Fisher Scientific, Cat# 047347.9M) in 1x PBS for 10 mins at room temperature while shaking, then washed and kept at 4 °C in PBS + 0.02% Na-azide until use.

#### Lentivirus packaging and lentiviral transduction

HEK293 T cells were maintained in Dulbecco’s Modified Eagles Medium (DMEM) high glucose with stable glutamine and sodium pyruvate (Capricorn Scientific) supplemented with 10% fetal bovine serum (GIBCO) and 1% Penicillin-Streptomycin (GIBCO). For POTATOMap proteomics, lentiviruses were produced by transient transfection of HEK293T cells with transfer vectors containing Streptavidin-KDEL-mNG or SBP-NLGN1-V5-APEX2, packaging vector psPAX2 and envelope vector pMD2.G with a ratio of 4:2:1 using PEI Max (Polysciences) according to standard protocols with a 3:1 PEI Max:DNA ratio for 6 to 9 h. PEI Max/DNA was mixed in fresh serum-free Opti-MEM medium (GIBCO), incubated for 20 min and added to the cells. All medium was removed after 6–9 h and changed to NB supplemented with 0.5 mM glutamine. Supernatants from packaging cells were collected 72 h post-transfection, filtered through a 0.45 µm filter and concentrated with Ultra Centrifugal Filter 100 kDa MWCO (AMICON) at 4 °C 1,000 g for 45 min. Lentivirus with Streptavidin-KDEL-mNG was made in batches and stored at -80 °C. Lentivirus with SBP-NLGN1-V5-APEX2 was immediately used to transduce DIV4 cortical neurons in RUSH medium. Cells were maintained at 37 °C and 5% CO2 until further processing at DIV8. For shRNA validation, lentiviruses were produced by transient transfection of HEK293T cells as described above, with transfer vector pLKO.1 puro containing shRNA, packaging vector psPAX2 and envelope vector pMD2.G with a ratio of 2.5:2:1 using PEI Max. Concentrated virus was stored in 4 °C until transduction of DIV4 cortical neurons in their original medium.

### Method details

#### DNA and shRNA constructs

The following vectors were used: pOC1 FLAG-SpCas9 (Addgene #183423), pCamK Homer1c-mRuby3, ptdTomato-C1, pJet1.2 Halotag donor, pX333 DRS-1-only FLAG-Cas9, ORANGEtrap destination vector, 3xalfa donor vector and pFSW_67 were a gift from Harold MacGillavry. pcDNA3_HA-KLC-TPR6 was a gift from Kristen Verhey^34^. TFR-GFP and HA-GLUR2 were gifts from Casper Hoogenraad. Halo-TFR-RUSH (Addgene #166905^85^) and PEGFP(A206K)-C1 were gifts from Jennifer Lippincott-Schwartz,. pNICE-NL2A was a gift from Peter Scheiffele (Addgene #15259^86^). psPAX2 (Addgene #12260) and PMD2.G (Addgene; #12259) were gifts from Didier Trono. FUGW (Addgene #14883^83^) was a gift from David Baltimore. pLKO.1 puro (Addgene #8453^84^) was a gift from Bob Weinberg.

Plasmids generated for this study:

FUGW_Strep-KDEL_SBP-NLGN1-V5-APEX2 was generated by amplifying the signal peptide and the rest of the protein (splice variant 4 or NLGN1B) separately from rat cDNA. SBP and V5-APEX2 were amplified from FUGW_Strep-KDEL_SBP-LAMP1-V5-APEX2 (Addgene #228524^18^). All fragments were inserted into FUGW_Strep-KDEL^18^ digested with EcoRI and BamHI by GIBSON assembly.

The following plasmids were generated by insertion of fragments into FUGW vector (Addgene plasmid # 14883) digested with XbaI and EcoRI by GIBSON assembly. FUGW_Strep-KDEL-mNG was generated by amplifying Strep-KDEL and mNG from FUGW_Strep-KDEL_SBP-SYT1-mNG^18^. FUGW_SBP-NLGN1-V5-APEX2 was generated by amplifying SBP-NLGN1-V5-APEX2 from FUGW_Strep-KDEL_SBP-NLGN1-V5-APEX2. FUGW_Strep-KDEL_mNG-V5-TFR-SBP was generated by amplifying mNG and SBP from FUGW_Strep-KDEL_SBP-SYT1-mNG, V5 from FUGW_Strep-KDEL_SBP-LAMP1-V5-APEX2 and TFR from Halo-TFR-SBP.

CMV_Strep-KDEL_SBP-NLGN1-Halo-FKBP was generated by amplifying Strep-KDEL-IRES-SBP-NLGN1 from FUGW_Strep-KDEL_SBP-NLGN1-V5-APEX2, Halotag was amplified from Halo-TFR-RUSH. 2xFKBP was amplified from mCh-FKBP2x-RTN4^89^, these were assembled into pEGFP-C1 (gift from Jennifer Lippincott-Schwartz) in between NheI and SacI restriction sites using GIBSON assembly.

mCherry-KIF5A-FRB was generated by amplifying FRB from GFP-BICD-N-FRB (gift from Lukas Kapitein) and KIF5A from mCherry-KIF5A-MD-Strep^87^, which were inserted into mCh-C1 inbetween XhoI and BamHI restriction sites using GIBSON assembly. HA-KIF5A-MD-FRB was generated by digesting mCherry-KIF5A-FRB with AgeI and BsrGI to remove mCherry and replacing it with annealed oligos encoding 3xHA

For subsequent cloning purposes, we generated the pFSW-StrepKDEL vector by amplification of Strep-KDEL-intron-IRES from Str-KDEL_SBP-EGFP-Ecadherin (Addgene plasmid # 65286) with XbaI and EcoRI sites introduced and inserted into restricted pFSW-67 (gift from Harold MacGillavry) by GIBSON assembly.

The following plasmids were generated by inserting their fragments into pFSW-StrepKDEL digested by EcoRI and BamHI using GIBSON assembly: pFSW_Strep-KDEL_SBP-NLGN1-mNG was generated by amplification of SBP-NLGN1 from FUGW_Strep-KDEL_SBP-NLGN1-V5-APEX2 and mNG from FUGW_Strep-KDEL_SBP-SYT1-mNG. pFSW_StrepKDEL_SBP-HA-NLGN1 was generated by amplification of SBP from FUGW_Strep-KDEL_SBP-NLGN1-V5-APEX2, and adding the HA tag by overhang PCR. NLGN1 was also amplified from FUGW_Strep-KDEL_SBP-NLGN1-V5-APEX2. pFSW_StrepKDEL_SBP-HA-mNG-NLGN1 was generated by amplification of SBP-HA from pFSW_StrepKDEL_SBP-HA-Nlgn1, mNG from FUGW_Strep-KDEL_SBP-SYT1-mNG and NLGN1 from FUGW_Strep-KDEL_SBP-NLGN1-V5-APEX2. pFSW_Str-KDEL_SBP-EGFP-HA-NLGN2 was generated by amplification of HA-NLGN2 from pNice_HA-NLGN2A (Addgene #15259). SBP-EGFP was amplified from pFSW_StrepKDEL_SBP-EGFP-NLGN1. pFSW_StrepKDEL_SBP-HA-NLGN1ΔWW and pFSW_StrepKDEL_SBP-HA-NLGN1ΔPDZ were generated by amplification from pFSW_StrepKDEL_SBP-HA-NLGN1. pFSW StrepKDEL_TfR-EGFP-SBP was generated by amplification of TFR from Halo-TFR-SBP, EGFP from PEGFP(A206K)-C1 and SBP from FUGW_Strep-KDEL_SBP-NLGN1-V5-APEX2. pFSW_StrKDEL_SBP-EGFP-HA-NLGN2ec-NLGN1ic was generated by amplification of SBP-EGFP-HA-NLGN2ec from pFSW_Str-KDEL_SBP-EGFP-HA-NLGN2 and NLGN1ic from FUGW_Strep-KDEL_SBP-NLGN1-V5-APEX2. pFSW_StrKDEL_SBP-EGFP-NLGN1ec-NLGN2ic was generated by amplification of SBP-EGFP-NLGN1ec from pFSW_Str-KDEL_SBP-EGFP-NLGN1 and NLGN2ic from pNice_HA-NLGN2A.

pFSW_StrepKDEL_SBP-EGFP-NLGN1 was generated by amplification of EGFP from PEGFP(A206K)-C1 and ligating this into pFSW_StrepKDEL_SBP-HA-Nlgn1 digested with BmtI and BamHI. β-actin_TENM2-EGFP was generated by amplification of TENM2 from rat cDNA in 3 fragments and inserting these into TFR-GFP digested with SalI and BmtI by GIBSON assembly.

To generate the NLGN1 knock-in construct (pX333_DRS-1-only FLAG-Cas9 NLGN1) we applied the HiUGE knock-in system^43^. The target sequence (CCACGTACTCTCTCAAAAGT) was selected using CRISPOR^90^ and incorporated into oligos, which were annealed and ligated into pX333 DRS-1-only FLAG-Cas9 which was digested with BpiI (gift from Harold MacGillavry).

To generate the NLGN1-smHA knock-in construct (pAAV_pOC2_NLGN1-ORANGE_smHA) we applied the pORANGEtrap system developed by the MacGillavry group (unpublished), which is based on the ORANGE system ^44,45^. ORANGEtrap destination vector and smHA donor plasmid with appropriate frameshift correction were gifts from Harold MacGillavry. The target sequence (TCTCTCATCTGGCTATACCC) was selected using CRISPOR^90^. Golden Gate cloning with Sap1 restriction enzyme (Thermofisher) was used to incorporate the annealed oligos and the 3xalfa donor sequence into the ORANGEtrap destination vector, in a similar fashion as the SapTrap method^91^. Similarly, we generated the NLGN2 knock-in construct (pAAV_pOC2_NLGN2-ORANGE_3xalfa) using ORANGEtrap destination vector and 3xalfa donor plasmid with appropriate frameshift correction (gifts from Harold MacGillavry). The target sequence used for NLGN2 was as following: GGGGCTGGCTGGGGCTCAACG.

The following sequences for rat-shRNAs inserted to pLKO.1-puro^84^ were used in this study: scramble shRNA (5’-GATGAAATATTCCGCAAGTAA-3’) was used for all experiments. KIF1A-shRNA (5’-CACGCCGTCTTCAACATCA-3’) was previously validated ^18,89^. The target sequences for other shRNAs used in this study are as follows: AFTPH (5’-CTTATCCATGGATTCTATTAA-3’), NSG2 (5’-TTGTTCCCATGTAAGATATTT-3’), MAGI3 (5’-TTACAGGAAAGGTCAAATATT-3’), APBB1(5’-GCCAATGCCAAGTGGTTAAAG-3’). These shRNAs have been validated in this study by quantitative PCR.

#### Antibodies and reagents

The following primary antibodies were used in this study: rabbit anti-TRIM46 (homemade, van Beuningen et al.^81^, 1:1000), mouse anti-V5 (Thermo Fisher Scientific Cat# R960-25, RRID:AB_2556564, 1:1000), rat anti-HA (Roche Cat# 11867423001, RRID:AB_390918, 1:200), mouse anti-pan-Neurofascin external (UC Davis/NIH NeuroMab Cat# 75-172, RRID:AB_2282826, 0.18 mg/ml), rabbit anti-GM130 (Abcam, RRID:AB_880266, 1:800), rabbit anti-ALFA (rHcAb) (NanoTag Biotechnologies Cat# N1583, RRID:AB_3075998, 1:1000), guinea pig anti-PSD95 (Kim et al., 1:2000), mouse anti-GABAA (Millipore Cat# MAB341, RRID:AB_11214320, 1:500), rabbit anti-Halo (Promega Cat# G9281, RRID:AB_713650, 1:500), mouse anti-GM130 (BD Biosciences Cat# 610823, RRID:AB_398142, 1:500), chicken anti-GFP (Aves Labs Cat# GFP-1020, RRID:AB_2307313, 1:500), chicken anti-VGAT (Synaptic Systems Cat# 131009, RRID:AB_2943610, 1:2000), mouse anti-gephyrin (Synaptic Systems Cat# 147011, RRID:AB_887717, 1:400), and mouse anti-AnkyrinG (Thermo Fisher Scientific Cat# 33-8800, RRID:AB_2533145, 1:200).

The following secondary antibodies were used in this study: goat anti-mouse Alexa Fluor 405 (Thermo Fisher Scientific Cat# A-31553, RRID:AB_221604, 1:500), goat anti-mouse IgG1 Alexa Fluor 647 (Thermo Fisher Scientific Cat# A21240, RRID:AB_2535809, 1:1000), goat anti-rabbit Alexa Fluor 405 (Thermo Fisher Scientific Cat# A-31556, RRID:AB_221605, 1:500), rabbit anti-mouse immunoglobulins/HRP (Agilent Cat# P0260, RRID:AB_2636929, 1:1000), goat anti-mouse IgG2a Alexa Fluor 594 (Thermo Fisher Scientific Cat# A-21135, RRID:AB_2535774, 1:1000), goat anti-rat Alexa Fluor 488 (Thermo Fisher Scientific Cat# A-11006, RRID:AB_2534074, 1:1000), goat anti-rat Alexa Fluor 568 (Thermo Fisher Scientific Cat# A-11077, RRID:AB_2534121, 1:1000), goat anti-rabbit Alexa Fluor 488 (Thermo Fisher Scientific Cat# A-11034, RRID:AB_2576217, 1:1000), goat anti-chicken Alexa Fluor 405 (Abcam Cat# ab175675, RRID:AB_2810980, 1:500), goat anti-chicken Alexa Fluor 568 (Thermo Fisher Scientific Cat# A11041, RRID:AB_2534098, 1:1000), and goat anti-guinea pig Alexa Fluor 647 (Thermo Fisher Scientific Cat# A-21450, RRID:AB_2535867, 1:1000).

The following primary antibodies were used for organotypic slice culturing: mouse anti-NLGN1 (SantaCruz Cat# sc-365110, RRID: AB_10708272), mouse anti-NLGN2 (Synaptic Systems Cat#129511, RRID: AB_2619813) rabbit anti-βIV-Spectrin (homemade, epitope corresponding to amino acids 2237-2256 of human βIV-Spectrin), guinea pig anti-AnkyrinG (Synaptic Systems Cat# 386004, RRID:AB_2725774), rabbit anti-vGat (Merck-Millipore, Cat#AB5062P, RRID:AB_2301998), chicken anti-Gephyrin (Synaptic Systems, Cat#147009; RRID:AB_2943527).

The following secondary antibodies were used for organotypic slice culturing: goat anti-chicken IgY AlexaFluor 405 (Abcam Cat #ab175675, RRID: AB_2810980), goat anti-mouse AlexaFluor 488 (Thermo Fisher Scientific Cat# A32723, RRID: AB_2633275), goat anti-rabbit AlexaFluor 568 (Thermo Fisher Scientific Cat# A11036, RRID: AB_10563566), goat anti-guinea pig Alexa Fluor 568 (Thermo Fisher Scientific Cat# A11075, RRID: AB_141954), goat anti-rabbit Alexa Fluor 647 (Thermo Fisher Scientific Cat# A21245, RRID: AB_2535813), goat anti-guinea pig AlexaFluor 647 (Thermo Fisher Scientific Cat# A21450, RRID: AB_2535867).

Other reagents used in this study were: Streptavidin Alexa Fluor 555 (Thermo Fisher Scientific Cat# S21381, RRID:AB_2307336, 1:500), FluoTag®-X2 anti-ALFA AZDye568 (NanoTag Biotechnologies Cat#N1502), biotin-phenol (Iris Biotech Cat#LS.3500), hydrogen peroxide (Sigma-Aldrich Cat#H1009), Lipofectamine 2000 (Thermo Fisher Scientific Cat#11668019), polyethylenimine (PEI MAX; Polysciences Cat#24765), Fluoromount-G mounting medium (Thermo Fisher Scientific Cat#00-4958-02), Pierce Streptavidin magnetic beads (Thermo Fisher Scientific Cat#88816), Trolox (Sigma-Aldrich Cat#238831), sodium L-ascorbate (Sigma-Aldrich Cat#0000374819), sodium azide (Sigma-Aldrich Cat#K43547688), TCEP (Sigma-Aldrich), iodoacetamide (IAA; Sigma-Aldrich), dithiothreitol (DTT; Sigma-Aldrich), bis-acrylamide 40% (Bio-Rad Cat#1610144), TEMED (Bio-Rad Cat#1610800), APS (Sigma-Aldrich Cat#A3678), anhydrous DMSO (Thermo Fisher Scientific Cat# D12345), Triton X-100 (Sigma-Aldrich Cat#93433), protease inhibitor cocktail (Roche), Rapalog (A/C heterodimerizer; Takara Bio Cat#635057), biotin (Sigma-Aldrich Cat#B4501), Dynasore (Tocris, 2897), Mix-n-Stain CF640R antibody labeling kit (Biotium Cat#92245), and JFX554 (Lavid Lab, Janelia).

#### RNA extraction, reverse transcription and quantitative PCR

We used cortical primary neurons to validate knockdown efficiency of AFTPH, NSG2, MAGI3 and APBB1 shRNA. Total RNA from transduced rat cortical neurons was extracted using RNeasy Mini kit (Qiagen, 74104). cDNA was reverse-transcribed using SuperScript III First-Strand Synthesis System (Invitrogen, 18080-051) according to manufacturer’s instructions. Quantification of target genes was done by qPCR using Power SYBR Green PCR Master Mix (Applied Biosystems, 4367659) and ViiA 7 Real-Time PCR System (Applied Biosystems). Data was extracted from ViiaA 7 Real-Time PCR system using QuantStudio Software V1.3 (Applied Biosystems). The following primers were used: APBB1 (GCTAAACCCGTTGGGGTAGA + GGTCCACTGCTCACGACTAC), MAGI3 (GGCCGGGTCATAGATGGAAG + GGTGGACCATGATGCTCTTCT), AFTPH (ATGGAACTCTGGACACCCTG + TTCAGTAAAGCTGTCTGAGCTG), NSG2 (TGCATTGTGTTCCTGGTGGT + GGATACAGCGCTTGTGCTTG).

#### Drug treatments

For the RUSH assay, neurons were incubated with 100 µM D-biotin and kept at 37 °C and 5% CO2 for the time indicated in each figure. For the axonal relocalization assay, neurons were incubated with 100 µM D-biotin at 37 °C and 5% CO2 for 45 min., then 1 µM rapalog or EtOH control was added and incubated at 37 °C and 5% CO2 for 3h 15 min. For Dynasore treatment together with RUSH release, 80 µM Dynasore and 100 µM D-biotin were added and incubated at 37 °C and 5% CO2 for 1h. For the AIS-network-activity assay, neurons were treated on DIV14 with either 10uM XE-991 or 2uM tetrodotoxin (TTX), 50uM AP5 and 40uM DNQX for 48 hours.

#### Immunofluorescence staining and confocal imaging

Neurons were incubated at room temperature with 4% paraformaldehyde supplemented with 4% sucrose in PBS for 5 min for fixation. Neurons were permeabilized with 0.2% Triton X-100 in PBS supplemented with calcium and magnesium (PBS-CM) for 10 min followed by blocking with 0.2% porcine gelatin in PBS-CM for 1 h at room temperature. Neurons were incubated with primary antibodies overnight at 4 °C or for 2 h at room temperature, washed three times with PBS-CM, incubated with secondary antibodies for 45 min at room temperature and washed three times with PBS-CM. Neurons were mounted in Fluoromount-G Mounting Medium (ThermoFisher Scientific). Neurons were imaged using a confocal laser-scanning microscope (LSM900, with Zen (blue edition) imaging software version 3.7.97.07000 (Zeiss)) equipped with Plan-Apochromat ×63 NA 1.40 oil DIC objective. Alternatively, neurons were imaged confocal laser-scanning microscope (LSM700, with Zen imaging software (Zeiss) version 8.1.7.484) equipped with Plan-Apochromat ×63 NA 1.40 oil DIC objective. For all IF experiments, hippocampal neurons were used, except for Fig. 1B, 1D and S1A.

For organotypic brain slice cultures, fixed slices were washed in PBS before being blocked for 2 h at room temperature in permeabilization solution (89.75% PBS, 10% normal goat serum, 0.25% Triton-X100 (from 10% Triton stock in PBS), while shaking. Antibodies were diluted in staining buffer (94.875% PBS, 5% normal goat serum, and 0.125% Triton X-100) and slices were incubated in this solution overnight at room temperature while shaking. Following three washes with PBS the next day, slices were incubated in secondary antibodies diluted in staining buffer for two hours at room temperature while shaking and under light exclusion. Stained slices were mounted onto coverslips using Vectashield (Vector Laboratories, Cat# H-1000-10) and then imaged using a confocal laser-scanning microscope (SP-8, Leica Microsystems). Images were acquired using a 1.4 NA 63x oil immersion lens at or approaching Nyquist resolution in X, Y, and Z.

#### Live-cell imaging

For live-cell imaging experiments, an inverted microscope Nikon Eclipse Ti-E (Nikon), equipped with a Plan Apo VC ×100 NA 1.40 oil objective (Nikon), a Yokogawa CSU-X1-A1 spinning disk confocal unit (Roper Scientific), a Photometrics Evolve 512 EMCCD camera (Roper Scientific) and an incubation chamber (Tokai Hit) mounted on a motorized XYZ stage (Applied Scientific Instrumentation) was used. MetaMorph (Molecular Devices) version 7.10.2.240 software was installed for controlling all devices. Coverslips mounted in a metal ring and supplemented in the original medium from hippocampal neurons were imaged in an incubation chamber that maintains optimal temperature and CO2 (37 °C and 5% CO2). To visualize multiple fluorescently labelled proteins and probes, sequential imaging was used, and each laser channel was exposed for 100—200 ms. Neurons were imaged every 1 s for 180 s. To identify the AIS, hippocampal neurons were incubated with a CF640R-conjugated antibody against neurofascin (NF-640R) for 10 min before live-cell imaging. To label HaloTag, JFX554 was incubated (100 nM in RUSH medium) for 10 min prior to biotin release of RUSH cargoes and returned to original RUSH medium.

#### Protein Origin, Trafficking And Targeting to Organelle Map (POTATOMap) sample preparation

DIV8 cortical neurons transduced with lentivirus at DIV4 stably expressing Strep-mNG-KDEL and SBP-NLGN1-V5-APEX2 (termed POTATOMap) were incubated with RUSH media supplemented with 500 µM biotin-phenol (IrisBio-tech) at 37 °C for 20 min before addition of 1 mM H2O2 (Sigma) at room temperature for 1 min to trigger peroxidase activity. Biotinylation was immediately quenched by two washes in ice-cold quencher solution (10 mM sodium azide, 10 mM sodium ascorbate and 5 mM Trolox in HBSS) and incubated 10 min on ice in azide-free quencher solution. Cells were scrapped and pellets were stored at −80 °C until all samples were harvested. Immunofluorescence was performed in the same batch of cortical neurons culture grown on 12-well plate to validate transduction efficiency. Cell pellets were lysed with freshly made quenching-RIPA buffer (150 mM NaCl, 50 mM Tris-HCl pH7.4, 0.1% SDS, 0.5% sodium-deoxycholate, 1% Triton X-100, 10 mM sodium azide, 10 mM sodium ascorbate, 5 mM Trolox and 1× protease inhibitors (Roche)) on ice for 30 min. Lysates were cleared by centrifuging at 16,000 × g 4 °C, supernatants were incubated overnight with pre-equilibrated Pierce magnetic streptavidin beads (Invitrogen, Cat#88817) on a rotor at 4 °C. Beads were washed three times with detergent-free quenching-RIPA buffer (150 mM NaCl, 50 mM Tris-HCl pH7.4, 10 mM sodium azide, 10 mM sodium ascorbate, 5 mM Trolox), three times freshly prepared Urea buffer (3M Urea in 50 mM ammonium bicarbonate) and three times freshly prepared 50mM ammonium bicarbonate solution (ABC). Beads were resuspended in 50 µL of 50mM ABC and reduced with 5 mM TCEP (Sigma) at 37°C for 30 min, alkylated with 10 mM IAA (Sigma) in the dark at room temperature for 30 min and quenched with 20 mM DTT (Sigma). Samples were washed two times with 50mM ABC and resuspended in 100 µL of the same buffer. Suspended beads were first incubated with 1 µg LysC for 4 hours at room temperature, followed by adding 1 µg Trypsin for digestion at 37 °C overnight. Peptides were collected by combining digested supernatant with two subsequent 100 µL 50mM ABC washes and immediately acidified with 1% trifluoroacetic acid. Digested peptides were desalted on Sep-Pak C18 Cartridges (Waters) and vacuum concentrated for storage until subsequent MS analysis.

#### Immunoblotting

Lysates were prepared as described above. Before bead incubation, 5% of lysates were taken for immunoblotting. Protein lysates were resolved by SDS-PAGE on a 10% Bis-Acrylamide (Bio-Rad) gel and transferred to a PVDF membrane (Bio-Rad). Membranes were blocked in 5% skimmed milk in TBS-T and washed three times with TBS-T. Then, they were incubated overnight at 4 °C with primary antibody (mouse anti-V5) in antibody buffer (3% BSA in TBS-T). After three washes with TBS-T, membranes were incubated with anti-mouse HRP secondary antibody in antibody buffer for 45 min at room temperature and again washed three times with TBS-T. Membranes were incubated in Clarity Western ECL Substrate (Bio-Rad, Cat#1705060) and developed using ImageQuant 800 (AMERSHAM). To visualize biotinylation, membranes were washed three times in TBS-T and incubated with IRDye 800CW Streptavidin for 45 min at room temperature. Membranes were washed two times in TBS-T and once in TBS and developed on an Odyssey CLx imaging system (LICOR) with Image Studio version 5.2.

#### MS and data analysis

All samples were reconstituted in 2% formic acid (FA) and analyzed on a Orbitrap Exploris 480 mass spectrometer (Thermo Fisher Scientific, San Jose, CA, United States) coupled to an UltiMate 3000 UHPLC system (Thermo Fisher Scientific, San Jose, CA, United States). Peptides were loaded onto a trap column (C18 PepMap100, 5 μm, 100 Å, 5 mm × 300 μm, Thermo Fisher Scientific, San Jose, CA, United States) with solvent A (0.1% formic acid in water) at 30 µl/min flowrate and chromatographically separated over the analytical column (Poroshell 120 EC C18, Agilent Technologies, 50 μm × 75 cm, 2.7 μm) using 180 min gradient at 300 nL/min flow rate. The gradient proceeds as follows: 9% solvent B (0.1% FA in 80% acetonitrile, 20% water) for 1 min, 9–13% for 1 min, 13–44% for 155 min, 44–55% for 5 min, 55–99% for 5 min, 99% for 3 min, and finally the system equilibrated with 9% B for 10 min.

The mass spectrometers were used in a data-dependent mode, which automatically switched between MS and MS/MS. After a survey MS scan ranging from 375 to 1600 m/z with 14 s dynamic exclusion time, the most abundant peptides of 120 m/z or higher were subjected to high energy collision dissociation (HCD) for further fragmentation using a 1.4 m/z isolation window. MS spectra were acquired in high-resolution mode (R > 60,000), whereas MS2 was in high-sensitivity mode (R > 15,000) and 28% normalized collision energy.

LC-MS/MS raw files were processed using MaxQuant’s (Version 2.4.2.0)^92^. for protein identification and label-free quantification (LFQ) using the integrated Andromeda search engine in reversed decoy mode against rat reference proteome (Uniprot-FASTA, UP000002494, downloaded March 2023) with an FDR of 0.01 at both peptide and protein levels. Digestion parameters were set to specific digestion with trypsin with a maximum number of 2 missed cleavage sites and a minimum peptide length of 7. Oxidation of methionine and amino-terminal acetylation were set as variable and carbamidomethylation of cysteine as fixed modifications. The tolerance window was kept at default. Label-free quantification is applied (minimum ratio count set to 2), and a total of 2 biological replicates with 2 technical replicates were analyzed. The resulting protein group file was processed and analyzed using R (Version 4.5.1) with a custom R script. From here onwards raw intensities were used for data analysis due to the nature of streptavidin affinity pull-down. Briefly, common contaminants, reverse, site-specific identifications were filtered out. Each condition were then separated into background control groups (w/o H2O2) or experimental groups (with H2O2) of their respective timepoint. For background control groups, proteins with less than 1 valid value were excluded. We assumed that values were missing not at random, hence remaining proteins were subjected to QRILC imputation using the R package MsCoreUtils and median normalized using proDA. Experimental groups with less than 3 valid values were excluded. Resulting protein groups were subjected to BPCA imputation and median normalized assuming values were missing at random due to detection limit. Each experimental group was joined back to their respective background control. Common proteins shared between experimental group and their respective background control of the same timepoint were tested using unpaired two sample t-test (p adjusted value < 0.05, log2 fold change ≥1) and annotated with fold change direction (Above or Below) and whether statistical significance is met (True or False). Proteins that were unique to experimental group of the specific timepoint were joined back to tested common experimental group and unique proteins were annotated as not detected in background. Control group data were removed and resulting in data frames with filtered proteins and background annotations in their respective timepoints (20min, 1h and 4h). Filtered timepoints were joint using Protein ID. To further analyze the data, we assumed that values were not missing at random across time due to biological relevance of our protein of interest (POI) trafficked throughout the biosynthetic pathway, hence missing values were imputed using QRILC and annotated how many valid values were present (3 or 4 or imputed) for clearer data representation. Imputation was done for statistical analysis purposes. Each timepoint was then tested against each other (1 h/20 min; 4 h/20 min; 4 h/1 h) using One way ANOVA with Tukey HSD corrections (adjusted p value < 0.05). For each comparison, we set specific thresholding for the log2 fold change: 20 min = diff_1H-20min <= -2; diff_4H-20min <= -2. 1 h = diff_1h-20mi >= 0.7; diff_4H-1H <= -1. 4 h = diff_4H-20min >= 0.7; diff_4H-1H >= 1. Lastly, we cross-referenced our dataset with an existing AIS immunoproximity proteome dataset ^32^. Data were further cleaned up in accordance with recent UniProt Update (November 2025) on reviewed and unreviewed proteins. To identify the temporal signature of a particular protein, Venn-diagram was constructed by identifying all significantly changing proteins in their respective test.

Proteins that were significantly detected against background in just one time point are termed ‘Unique Proteins’. For each time point individually, these proteins were analyzed by STRING Gene Enrichment. A selection of enriched terms and the proteins in these terms are highlighted in Fig S1C.

EnrichGO package was used for Gene Ontology analysis in Fig. 1F. The background universe was set to the total number of proteins analyzed after filtering for contaminants (3608 proteins). All terms were filtered (padjust <0.05) and simplified. Dot plots were generated by plotting each selected term according to their enriched time point, number of proteins categorized in that term as size, and a color gradient was used to depict the -log10 padjust value (Fig. 1F).

Figure 1G was generated by grouping proteins within enriched terms. Within these groups, proteins were further segregated manually according to known function. For Golgi Apparatus, we depicted a selection of proteins. Proteins enriched within each of the 3 time points, were compared with our previously published LAMP1 POTATOMap dataset^18^ and a Venn-diagram was constructed to display the similarities (Fig S1E). Proteins shared between NLGN1 20 min. time point and LAMP1 1h time point were subjected to STRING Gene Enrichment analysis (Figure S1E).

For the identification of potential co-cargoes, we filtered all proteins for their presence at the 20 min. and 1h time points (columns Background_info_X are either: ‘ABOVE_TRUE’ or ‘Not Detected in Background’). Using Uniprot ID mapping, we filtered out the hits that have a transmembrane domain. These equal the 20min1h membrane set. Then, we analyzed this set of proteins by STRING Gene Enrichment and selected all hits mapped to the term ‘plasma membrane’, these form the 20min1h PM set. Next, we retrieved all postsynaptic proteins from the SynGO database ^93^. Using Uniprot ID mapping, we filtered out the proteins with a transmembrane domain. This set was further analyzed by STRING Gene Enrichment. We selected all hits in the term ‘PM’. These form the postsynaptic PM protein set.

Heat maps were generated by selecting specific hits and plotted according to their respective log2 Fold Change. Imputed or missing values in one of the time points are depicted in grey.

#### Image analysis and quantification

Fluorescence line intensity plots: co-distribution of different proteins was analyzed using Image J. Plot profiles were generated using a line traced along specified markers. Length of traced profile line is indicated in each intensity plot.

Kymograph analysis: Kymographs were generated from live cell images using Image J. Segmented lines of approximately 30 μm were drawn and straightened along the AIS identified using the AIS marker NF-CF640R. Straightened AISs were re-sliced followed by z-projection to obtain kymograph. A random segment of 30μm was cropped out for analysis. Anterograde movements were oriented from left to right in all kymographs. Time of recording and length of segments are indicated in each kymograph. The number of events for anterograde, retrograde, stationary and total number of RUSH-NLGN1 and RUSH-NLGN2 were obtained from three independent experiments. To measure velocity of trajectories (Fig. S4A), straight line ROIs were manually drawn along mobile segments of individual traces. The angle of the lines was used to calculate the average velocity of the mobile segments.

AIS/dendrite ratio was measured by tracing segments in 3 dendrites and the AIS. For each cell, the ratio was calculated by dividing the intensity at the AIS by the mean intensity at the dendrites.

Normalized intensity at the AIS was measured by tracing and measuring the intensity at the AIS and measuring the intensity in the soma. For each cell, the AIS intensity was normalized to the intensity of the soma.

Polarity Index was measured by tracing segments in 3 dendrites and the axon or AIS. For older neurons (DIV15), thickness of the line was set to 15 pixels. Polarity Index was calculated as: (mean intensity dendrites – mean intensity axon or AIS) / (mean intensity dendrites + mean intensity axon or AIS). In crosslinking experiments, the shift in polarity was calculated as the difference between the average polarity in control and crosslinked conditions within each replicate.

CellProfiler was used to analyze the distribution of endogenous NLGN1 and NLGN2. To select the AIS, Gaussian filter and RescaleIntensity were applied to the channel with the AIS marker. IdentifyPrimaryObjects was used to select all AIS’s in the field of view. For knock-in images, AIS of knock-in cells were identified by making a rough selection of NLGN puncta by IdentifyPrimaryObjects, followed by RelateObjects. For images stained for Neuroligin, we imaged field of views with containing just one AIS. We then expanded the AIS-object to mask puncta at the plasma membrane and presynapse. This mask was used to select NLGN1, NLGN2 and VGAT puncta, which were found by IdentifyPrimaryObjects. Masks of these objects were used to calculate the Mander’s colocalization. RelateObjects was just to find the fraction of puncta colocalizing with each other. The same pipeline was used to analyze the colocalization between NLGN and GEPH in Fig. 8G. To analyze NLGN2 intensity along the AIS, a line plot was drawn over the entire length of the AIS-object-masked NLGN2 images that were imported into FIJI/ImageJ. The measured intensity across AIS length was binned into bins of 10% AIS length. To analyze AIS length and distance from soma the ProFeatFit FIJI plugin was used.

To identify NLGN1/2 puncta on the AIS, we used the FIJI plugin SNT^94^ to trace the AIS and export a filled mask of these traces as binary images, with which we then masked the corresponding NLGN1/2 channel. In the FIJI plugin 3D Suite^95^, we then segmented both the AIS and AIS-filtered NLGN1/2 channels via Simple Segmentation and ran 3D Multi-Coloc to link AIS to AIS-borne NLGN puncta. For NLGN2 colocalization with synaptic structures, NLGN2 puncta were identified by eye, with channels for pre- and postsynapses turned off, noting down the corresponding position of the puncta (proximal/medial/distal). Afterwards, synaptic channels were turned back on and colocalization was assessed by eye. Line profiles were drawn for illustrative purposes. NLGN2 location index along the AIS was defined as the weighted average of NLGN2 puncta positions over total number of NLGN2 puncta for each AIS. Where applicable, we used the FIJI-native Bleach Correction feature set to histogram or exponential fit to account for axial fluorescence dropoff.

#### Statistical analysis

Data processing and statistical analysis were performed using Microsoft Excel, GraphPad Prism (version 9.5.1 and 11.0.1), MaxQuant (Version 2.0.1.0), Perseus (version 1.6.15.0), VScode and RStudio. Significance was determined as following: ns- not significant, *p < 0.05, **p < 0.01, ***p < 0.001, and ****p < 0.0001. Data normality was checked using D’Agostino–Pearson omnibus test. The statistical test performed, number of cells (n) and independent experiments (N) are indicated in figure legends. Exact p values are reported in figure legends and Source Data. No statistical method was used to predetermine sample size. The experiments were not randomized and investigators were not blinded to allocation during experiments and outcome assessment.

---

**Figure S1 (related to Fig. 1.).**
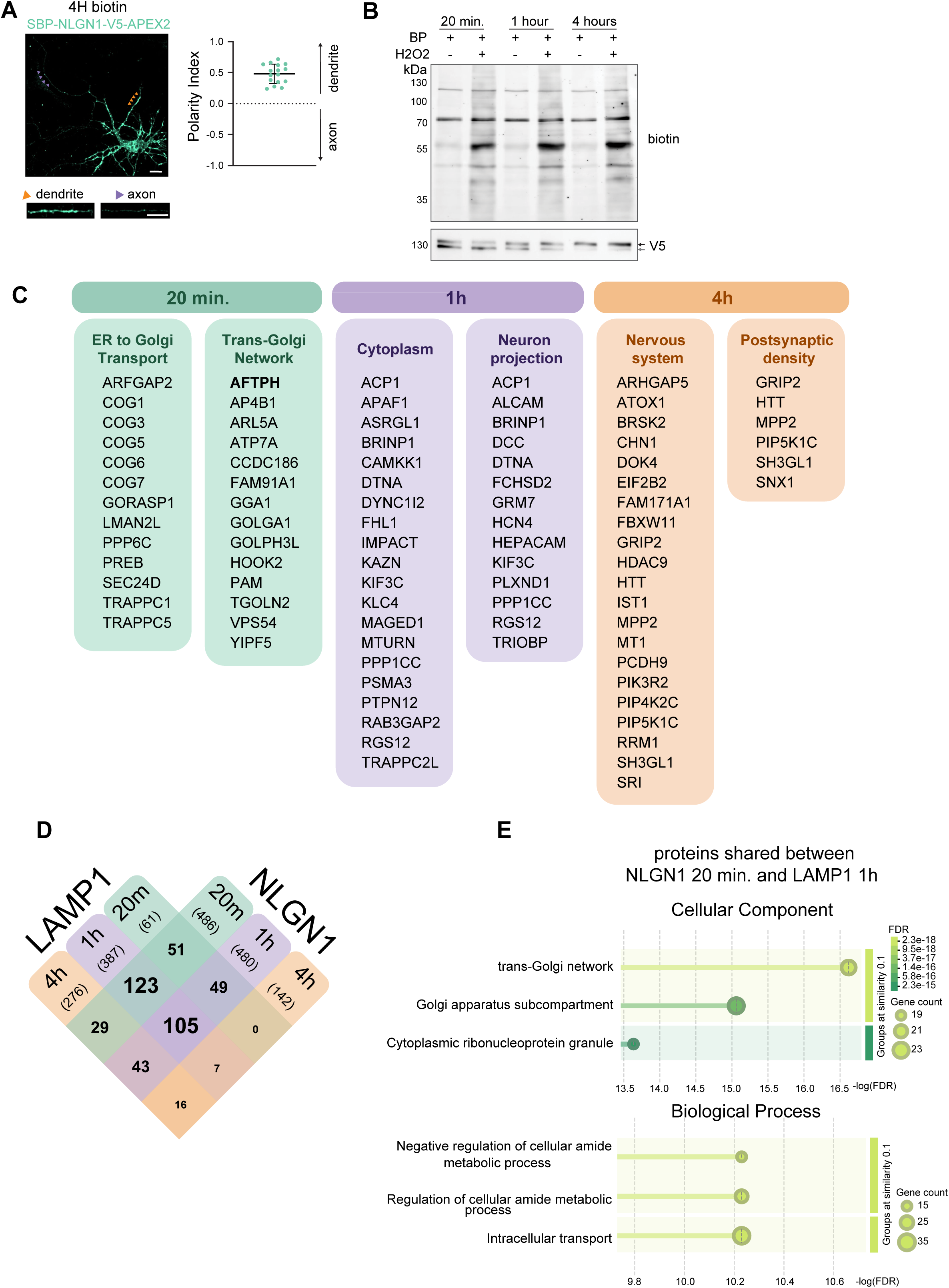
Spatiotemporal interactome of biosynthetic NLGN1 during its trafficking from ER to PM. (A) Representative image a SBP-NLGN1-V5-APEX2 and Strep-mNG-KDEL transduced cortical neuron, incubated with biotin for 4h, and stained for V5 (left). Graph shows the Polarity Index; each data point represents a neuron, N = 2 independent experiments (right). (B) Immunoblot of SBP-NLGN1-V5-APEX2 and Strep-mNG-KDEL transduced neurons lysed at 20min, 1h and 4h time points, with and without H2O2, stained for biotin with conjugated Streptavidin (top) and V5 (bottom). Arrows indicate immature (bottom) and mature (top) glycosylation states. (C) Schematic representation of Unique Proteins (proteins with 2 imputed values), grouped based on Gene Ontology. (D-E) Venn diagram comparing enriched proteins per time point between LAMP1 and NLGN1 (D). STRING Gene Ontology Enrichment analysis of proteins shared between LAMP1 20 min. and NLGN1 1h (E).

**Figure S2 (related to Fig. 2).**
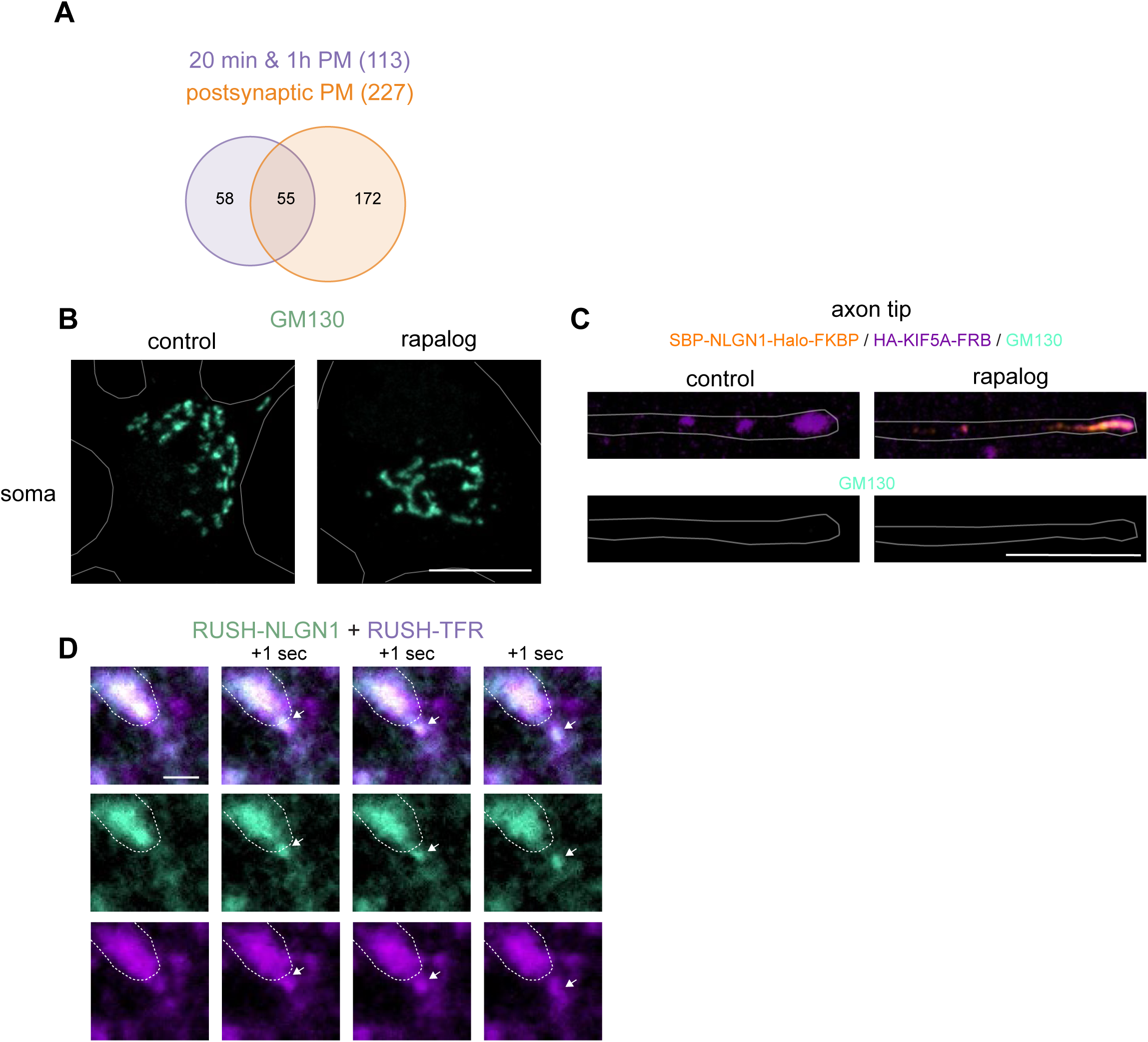
NLGN1 cotraffics with a subset of secreted cargoes. (A) Venn diagram of 20 min. & 1h PM proteins compared to postsynaptic PM proteins. (B-C) Representative images of soma (B) and axon tips (C) of hippocampal neurons expressing SBP-NLGN1-Halo-FKBP and HA-KIF5A-FRB, treated as shown in Fig 2D, with biotin and with EtOH (control) or Rapalog and stained for GM130. Scale bars represent 10 µm. (D) Still images of live-imaged DIV8 neurons transfected with RUSH-NLGN1-mNG and Halo-TFR-RUSH, incubated with biotin for 1h. Arrow indicates vesicle budding. Scale bar represents 1 µm.

**Figure S3 (related to Fig. 3).**
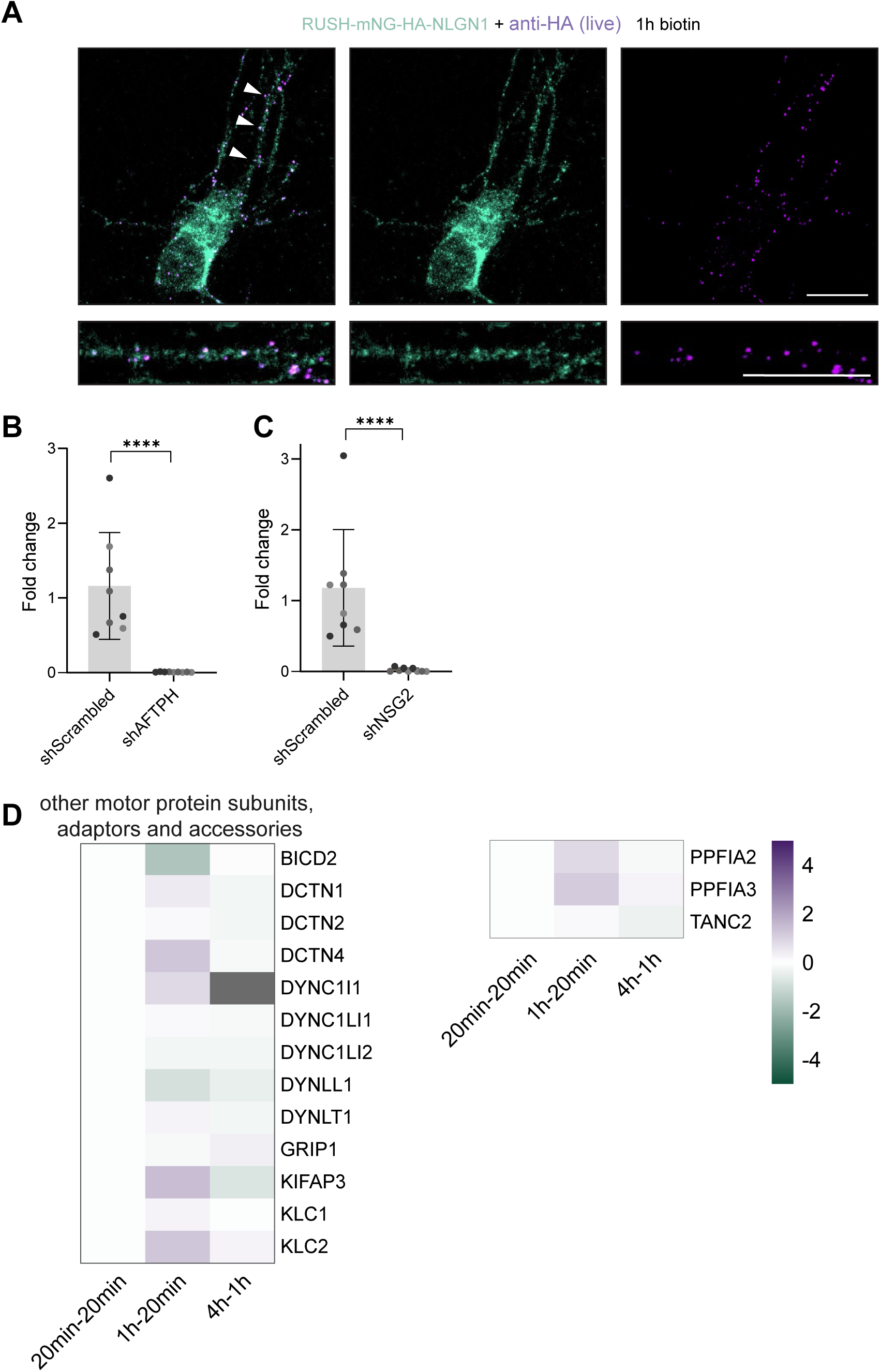
Validation of NLGN1 surface labeling and AFTPH and NSG2 knockdowns. (A) Representative overview image and dendrite crop of a neuron expressing SBP-mNG-HA-NLGN1, live incubated with biotin and surface labeled for HA for 1h. Secondary antibody was incubated after fixation, before permeabilization. Scale bar represents 10 µm. Arrows indicate dendrite. (B-C) Quantitative PCR fold change of shAFTPH (B) and shNSG2 (C) compared to shScrambled. All values were normalized to the respective GAPDH internal control. Unpaired t test with Welch’s correction (B); unpaired t test (C). Colors represent independent experiments (3), individual dots represent technical replicates. (D) Heat map of motor protein subunits, adaptors and accessories, in which the log2 Fold Change across time points is plotted.

**Figure S4 (related to Fig. 4).**
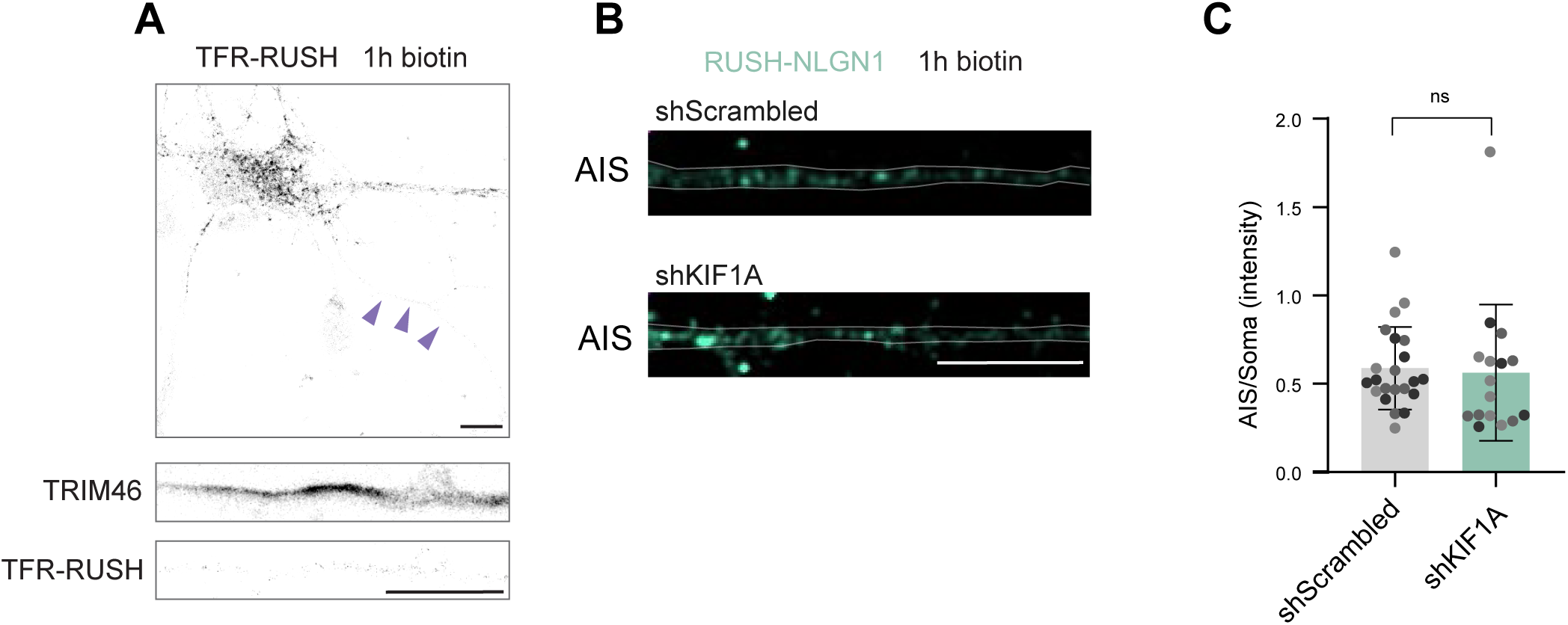
TFR-RUSH is not sorted to the AIS and kinesin-3 is not involved in the transport of RUSH-NLGN1 to the AIS. (A) Representative image of soma and AIS of a neuron expressing mNG-V5-TFR-RUSH, incubated with biotin for 1h, and stained for TRIM46. Arrows indicate the AIS. (B) Representative images of AIS of neuronsexpressing RUSH-HA-NLGN1, tdTomato (fill) and shScrambled or shKIF1A, incubated with biotin for 1h and surface labeled for HA. See also Fig. 3D. Quantification AIS/Soma intensity ratio, tested with Mann-Whitney test. Data points represent individual neurons, color indicates individual experiments. (C). Scale bars represent 10 µm.

**Figure S5 (related to Fig. 5).**
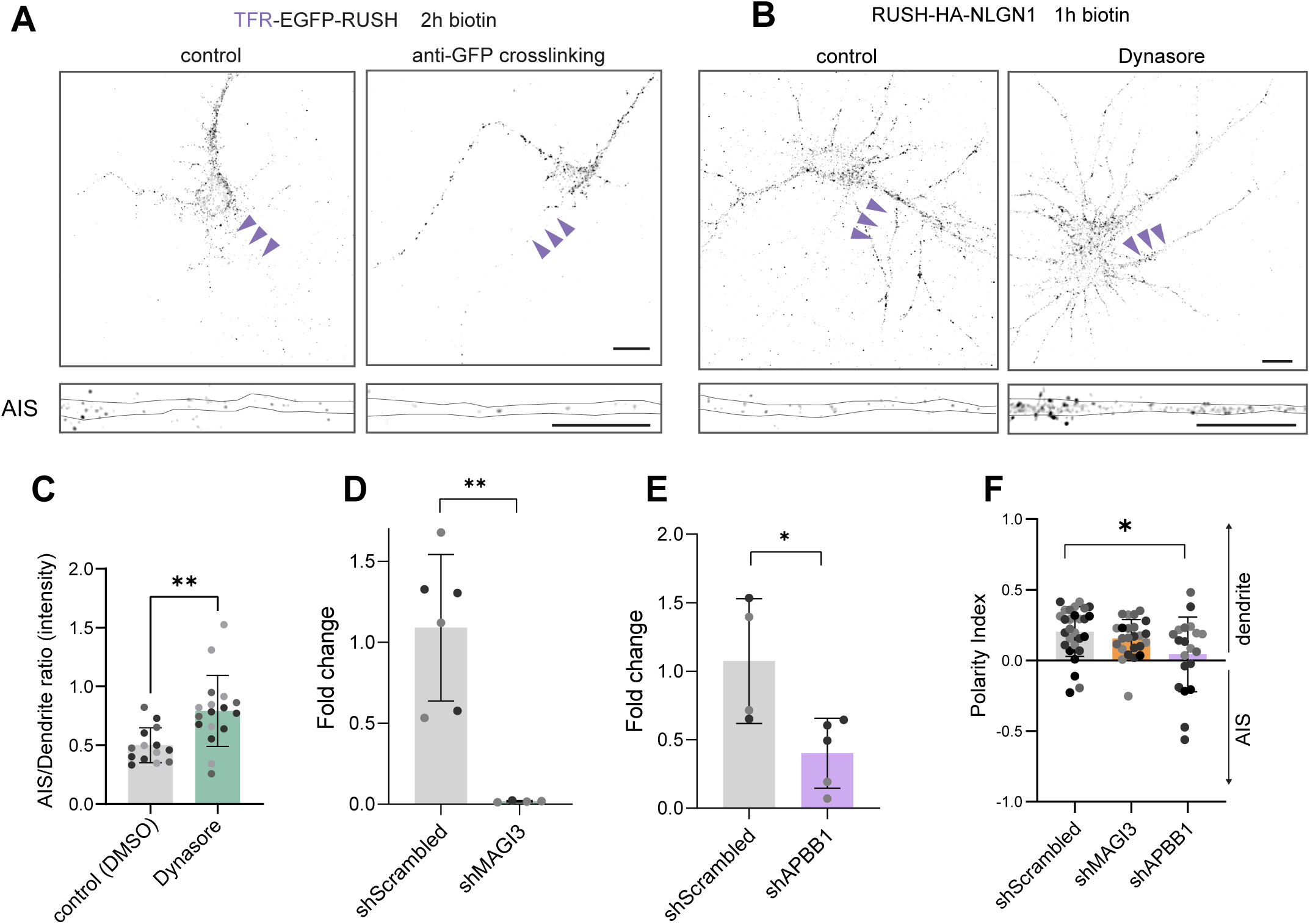
Removal of NLGN1 from the AIS PM is mediated by endocytosis, APBB1, MAGI3. (A) Representative images of soma and AIS of neurons expressing TFR-EGFP-RUSH, live incubated with biotin for 2h in the presence of anti-GFP antibody(crosslinking) or stained for GFP after fixation (control). (B-C) Representative images of soma and AIS of neurons expressing RUSH-HA-NLGN1, live-incubated with biotin for 1h, in the presence of DMSO (control) or 80 µM Dynasore (B), and surface labeled for HA. Quantification of AIS/dendrite intensity ratio, tested with Mann-Whitney test. Data points represent individual neurons, colors indicate the individual experiments (C). (D-E) Quantitative PCR fold change of shMAGI3 (D) or shAPBB1 (E) compared to shScrambled. All values were normalized to respective GAPDH internal control. Tested with unpaired t test (D, E). Colors represent independent experiments (2), individual dots represent technical replicates. (F) Quantification of Polarity Index with Kruskal-Wallis test with Dunn’s correction. Arrows indicate the AIS. Scale bars represent 10 µm.

**Figure S6 (related to Fig. 6).**
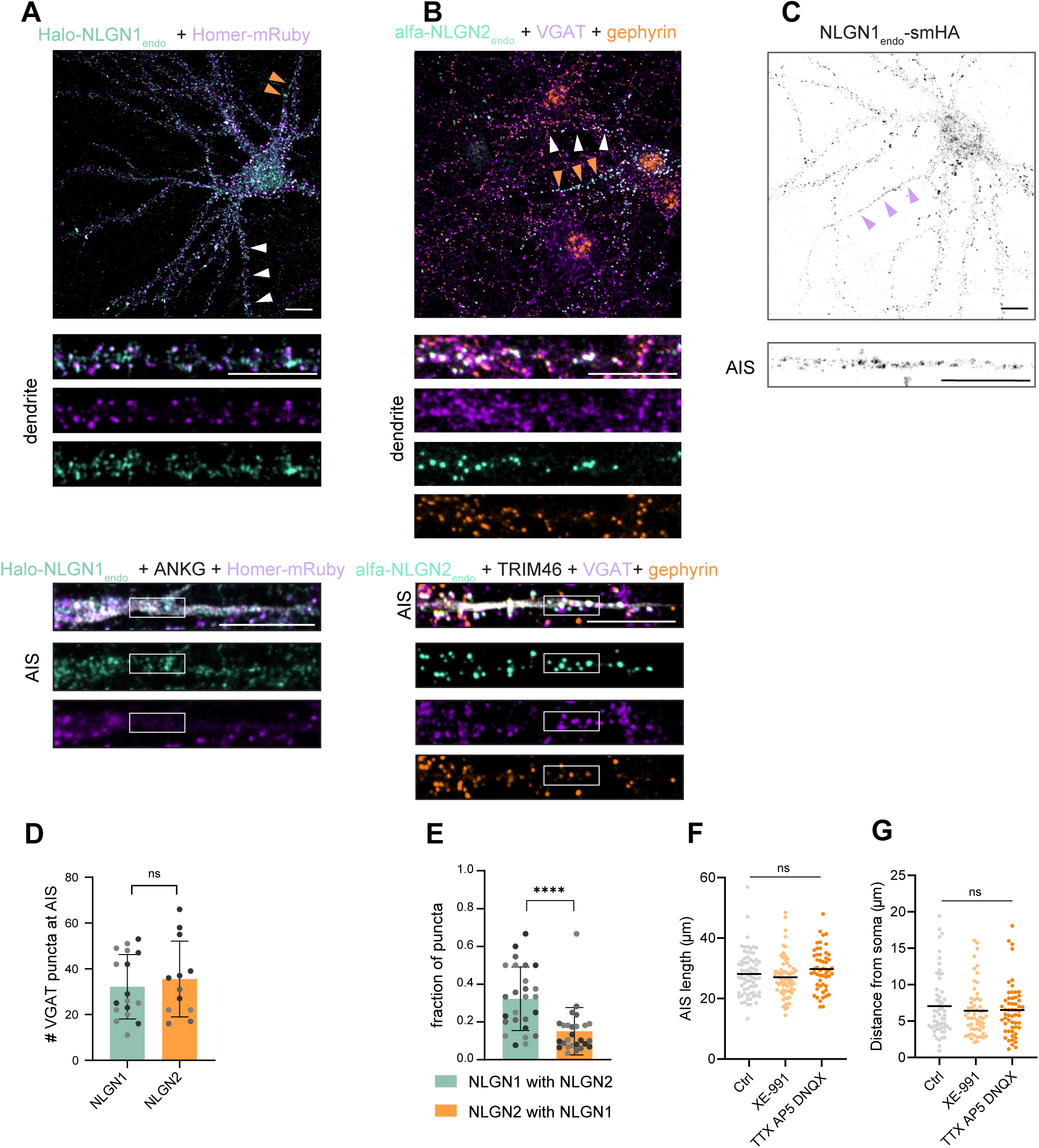
NLGN2 predominantly occupies inhibitory synapses at the AIS. (A-B) Representative overview images and dendrite and AIS crops of fully mature neurons (DIV21), with NLGN1 endogenously tagged with HaloTag at the N-terminus, expressing Homer1c-mRuby3 and stained for ANKG (A) and NLGN2 endogenously tagged with 3xalfa at the N-terminus (B), stained for Trim46, VGAT and GEPH (B). (C) Representative overview image and AIS crop of neuron with NLGN1 endogenously tagged with smHA at the C-terminus. (D) Quantification of number of VGAT puncta at the AIS, corresponding to Fig. 6A, B. unpaired t test. (E) Quantification of the fraction of NLGN1, NLGN2 puncta colocalizing with NLGN2 and NLGN1, respectively. Wilcoxon matched-pairs signed rank test. (F) Quantification of AIS length. Kruskal-Wallis test with Dunn’s correction, N=3 independent experiment, n=53-72 cells (G) Quantification of distance between AIS and soma. Kruskal-Wallis test with Dunn’s correction, N=3 experiments. N=3 independent experiments, n=56-57 cells. Data points represent individual neurons, colors indicate the individual experiments. Orange/purple arrows indicate the AIS, white arrows indicate the dendrite. Scale bars represent 10 µm.

**Figure S7 (related to Fig. 7).**
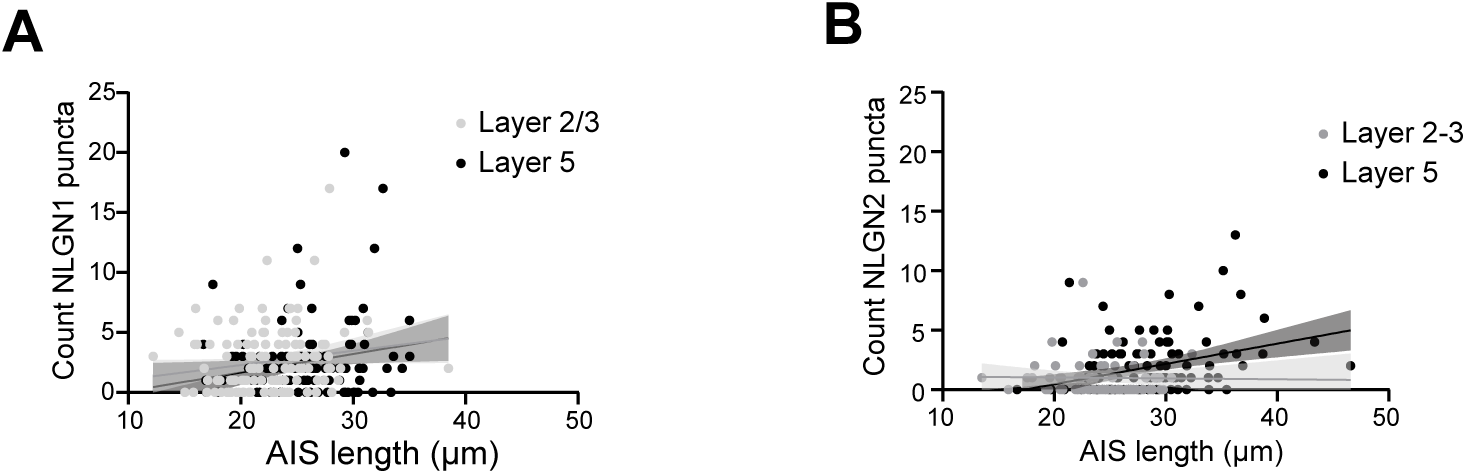
(A) Count of NLGN1 puncta plotted against the AIS length (R^2^ = 0.0305 and 0.0413 with P = 0.0576 and 0.0316 for L2/3). Fisher’s exact test. (B) Count of NLGN2 puncta plotted against the AIS length (R^2^ = 0.0127 and 0.1606, with P = 0.3144 and <0.0001 for L2/3 and L5). Fisher’s exact test.

## References

1. Matsuda, S., Miura, E., Matsuda, K., Kakegawa, W., Kohda, K., Watanabe, M., and Yuzaki, M. (2008). Accumulation of AMPA Receptors in Autophagosomes in Neuronal Axons Lacking Adaptor Protein AP-4. Neuron 57, 730–745. 10.1016/j.neuron.2008.02.012.

2. Farías, G.G., Cuitino, L., Guo, X., Ren, X., Jarnik, M., Mattera, R., and Bonifacino, J.S. (2012). Signal-Mediated, AP-1/Clathrin-Dependent Sorting of Transmembrane Receptors to the Somatodendritic Domain of Hippocampal Neurons. Neuron 75, 810–823. 10.1016/j.neuron.2012.07.007.

3. Li, P., Merrill, S.A., Jorgensen, E.M., and Shen, K. (2016). Two Clathrin Adaptor Protein Complexes Instruct Axon-Dendrite Polarity. Neuron 90, 564–580. 10.1016/j.neuron.2016.04.020.

4. Bentley, M., and Banker, G. (2016). The cellular mechanisms that maintain neuronal polarity. Nature Reviews Neuroscience 17, 611–622. 10.1038/nrn.2016.100.

5. Guardia, C.M., De Pace, R., Mattera, R., and Bonifacino, J.S. (2018). Neuronal functions of adaptor complexes involved in protein sorting. Current Opinion in Neurobiology 51, 103–110. 10.1016/j.conb.2018.02.021.

6. Kersten, N., and Farías, G.G. (2023). A voyage from the ER: spatiotemporal insights into polarized protein secretion in neurons. Front. Cell Dev. Biol. 11. 10.3389/fcell.2023.1333738.

7. Südhof, T.C. (2008). Neuroligins and neurexins link synaptic function to cognitive disease. Nature 455, 903–911. 10.1038/nature07456.

8. Bemben, M.A., Shipman, S.L., Nicoll, R.A., and Roche, K.W. (2015). The Cellular and Molecular Landscape of Neuroligins. Trends Neurosci 38, 496–505. 10.1016/j.tins.2015.06.004.

9. Han, Y., Dai, X., and Zhang, B. (2026). Orchestrating synaptic strength by neuroligin-confined nanoscale organization. Molecules and Cells 49, 100319. 10.1016/j.mocell.2026.100319.

10. Jamain, S., Quach, H., Betancur, C., Råstam, M., Colineaux, C., Gillberg, I.C., Soderstrom, H., Giros, B., Leboyer, M., Gillberg, C., et al. (2003). Mutations of the X-linked genes encoding neuroligins NLGN3 and NLGN4 are associated with autism. Nat Genet 34, 27–29. 10.1038/ng1136.

11. Sun, C., Cheng, M.-C., Qin, R., Liao, D.-L., Chen, T.-T., Koong, F.-J., Chen, G., and Chen, C.-H. (2011). Identification and functional characterization of rare mutations of the neuroligin-2 gene (NLGN2) associated with schizophrenia. Hum Mol Genet 20, 3042–3051. 10.1093/hmg/ddr208.

12. Cao, F., Liu, J.J., Zhou, S., Cortez, M.A., Snead, O.C., Han, J., and Jia, Z. (2020). Neuroligin 2 regulates absence seizures and behavioral arrests through GABAergic transmission within the thalamocortical circuitry. Nat Commun 11, 3744. 10.1038/s41467-020-17560-3.

13. Chih, B., Afridi, S.K., Clark, L., and Scheiffele, P. (2004). Disorder-associated mutations lead to functional inactivation of neuroligins. Hum Mol Genet 13, 1471–1477. 10.1093/hmg/ddh158.

14. Chih, B., Engelman, H., and Scheiffele, P. (2005). Control of Excitatory and Inhibitory Synapse Formation by Neuroligins. Science 307, 1324–1328. 10.1126/science.1107470.

15. Varoqueaux, F., Aramuni, G., Rawson, R.L., Mohrmann, R., Missler, M., Gottmann, K., Zhang, W., Südhof, T.C., and Brose, N. (2006). Neuroligins Determine Synapse Maturation and Function. Neuron 51, 741–754. 10.1016/j.neuron.2006.09.003.

16. Wang, J., Sudhof, T., and Wernig, M. (2025). Distinct mechanisms control the specific synaptic functions of Neuroligin 1 and Neuroligin 2. EMBO reports 26, 860–879. 10.1038/s44319-024-00286-4.

17. Bourke, A.M., Schwartz, S.L., Bowen, A.B., Kleinjan, M.S., Winborn, C.S., Kareemo, D.J., Gutnick, A., Schwarz, T.L., and Kennedy, M.J. (2021). zapERtrap: A light-regulated ER release system reveals unexpected neuronal trafficking pathways. Journal of Cell Biology 220. 10.1083/jcb.202103186.

18. Li, C.H., Kersten, N., Özkan, N., Nguyen, D.T.M., Koppers, M., Post, H., Altelaar, M., and Farias, G.G. (2024). Spatiotemporal proteomics reveals the biosynthetic lysosomal membrane protein interactome in neurons. Nat Commun 15, 10829. 10.1038/s41467-024-55052-w.

19. Boncompain, G., Divoux, S., Gareil, N., de Forges, H., Lescure, A., Latreche, L., Mercanti, V., Jollivet, F., Raposo, G., and Perez, F. (2012). Synchronization of secretory protein traffic in populations of cells. Nature Methods 9, 493–498. 10.1038/nmeth.1928.

20. Hung, V., Udeshi, N.D., Lam, S.S., Loh, K.H., Cox, K.J., Pedram, K., Carr, S.A., and Ting, A.Y. (2016). Spatially resolved proteomic mapping in living cells with the engineered peroxidase APEX2. Nature Protocols 11, 456–475. 10.1038/nprot.2016.018.

21. Mattera, R., Ritter, B., Sidhu, S.S., McPherson, P.S., and Bonifacino, J.S. (2004). Definition of the Consensus Motif Recognized by γ-Adaptin Ear Domains*. Journal of Biological Chemistry 279, 8018–8028. 10.1074/jbc.M311873200.

22. Burman, J.L., Wasiak, S., Ritter, B., de Heuvel, E., and McPherson, P.S. (2005). Aftiphilin is a component of the clathrin machinery in neurons. FEBS Letters 579, 2177–2184. 10.1016/j.febslet.2005.03.008.

23. Sabéran-Djoneidi, D., Marey-Semper, I., Picart, R., Studler, J.-M., Tougard, C., Glowinski, J., and Lévi-Strauss, M. (1995). A 19-kDa Protein Belonging to a New Family Is Expressed in the Golgi Apparatus of Neural Cells (*). Journal of Biological Chemistry 270, 1888–1893. 10.1074/jbc.270.4.1888.

24. Xiao, J., Dai, R., Negyessy, L., and Bergson, C. (2006). Calcyon, a Novel Partner of Clathrin Light Chain, Stimulates Clathrin-mediated Endocytosis*. Journal of Biological Chemistry 281, 15182–15193. 10.1074/jbc.M600265200.

25. Lo, K.Y., Kuzmin, A., Unger, S.M., Petersen, J.D., and Silverman, M.A. (2011). KIF1A is the primary anterograde motor protein required for the axonal transport of dense-core vesicles in cultured hippocampal neurons. Neuroscience Letters 491, 168–173. 10.1016/j.neulet.2011.01.018.

26. Stucchi, R., Plucińska, G., Hummel, J.J.A., Zahavi, E.E., Guerra San Juan, I., Klykov, O., Scheltema, R.A., Altelaar, A.F.M., and Hoogenraad, C.C. (2018). Regulation of KIF1A-Driven Dense Core Vesicle Transport: Ca2+/CaM Controls DCV Binding and Liprin-α/TANC2 Recruits DCVs to Postsynaptic Sites. Cell Reports 24, 685–700. 10.1016/j.celrep.2018.06.071.

27. Watanabe, K., Al-Bassam, S., Miyazaki, Y., Wandless, T.J., Webster, P., and Arnold, D.B. (2012). Networks of Polarized Actin Filaments in the Axon Initial Segment Provide a Mechanism for Sorting Axonal and Dendritic Proteins. Cell Reports 2, 1546–1553. 10.1016/j.celrep.2012.11.015.

28. Kuijpers, M., van de Willige, D., Freal, A., Chazeau, A., Franker, M.A., Hofenk, J., Rodrigues, R.J.C., Kapitein, L.C., Akhmanova, A., Jaarsma, D., et al. (2016). Dynein Regulator NDEL1 Controls Polarized Cargo Transport at the Axon Initial Segment. Neuron 89, 461–471. 10.1016/j.neuron.2016.01.022.

29. Balasanyan, V., Watanabe, K., Dempsey, W.P., Lewis, T.L., Trinh, L.A., and Arnold, D.B. (2017). Structure and Function of an Actin-Based Filter in the Proximal Axon. Cell Reports 21, 2696–2705. 10.1016/j.celrep.2017.11.046.

30. Jin Ye, Jin Ye, Ye, J., Li, J., Jianchao Li, Li, J., Fei Ye, Ye, F., Yan Zhang, Zhang, Y., et al. (2020). Mechanistic insights into the interactions of dynein regulator Ndel1 with neuronal ankyrins and implications in polarity maintenance. Proceedings of the National Academy of Sciences of the United States of America 117, 1207–1215. 10.1073/pnas.1916987117.

31. Farías, G.G., Guardia, C.M., Britt, D.J., Guo, X., and Bonifacino, J.S. (2015). Sorting of Dendritic and Axonal Vesicles at the Pre-axonal Exclusion Zone. Cell Rep 13, 1221–1232. 10.1016/j.celrep.2015.09.074.

32. Zhang, W., Fu, Y., Peng, L., Ogawa, Y., Ding, X., Rasband, A., Zhou, X., Shelly, M., Rasband, M.N., and Zou, P. (2023). Immunoproximity biotinylation reveals the axon initial segment proteome. Nat Commun 14, 8201. 10.1038/s41467-023-44015-2.

33. Zahavi, E.E., Hummel, J.J.A., Han, Y., Bar, C., Stucchi, R., Altelaar, M., and Hoogenraad, C.C. (2021). Combined kinesin-1 and kinesin-3 activity drives axonal trafficking of TrkB receptors in Rab6 carriers. Developmental Cell 56, 494–508.e7. 10.1016/j.devcel.2021.01.010.

34. Verhey, K.J., Meyer, D., Deehan, R., Blenis, J., Schnapp, B.J., Rapoport, T.A., and Margolis, B. (2001). Cargo of Kinesin Identified as Jip Scaffolding Proteins and Associated Signaling Molecules. J Cell Biol 152, 959–970. 10.1083/jcb.152.5.959.

35. Eichel, K., Uenaka, T., Belapurkar, V., Lu, R., Cheng, S., Pak, J.S., Taylor, C.A., Südhof, T.C., Malenka, R., Wernig, M., et al. (2022). Endocytosis in the axon initial segment maintains neuronal polarity. Nature 609, 128–135. 10.1038/s41586-022-05074-5.

36. Iida, J., Hirabayashi, S., Sato, Y., and Hata, Y. (2004). Synaptic scaffolding molecule is involved in the synaptic clustering of neuroligin. Molecular and Cellular Neuroscience 27, 497–508. 10.1016/j.mcn.2004.08.006.

37. Jeong, J., Pandey, S., Li, Y., John D. Badger, I.I., Lu, W., and Roche, K.W. (2019). PSD-95 binding dynamically regulates NLGN1 trafficking and function. Proceedings of the National Academy of Sciences of the United States of America 116, 12035. 10.1073/pnas.1821775116.

38. Jiang, S., Li, Y., Zhang, X., Bu, G., Xu, H., and Zhang, Y. (2014). Trafficking regulation of proteins in Alzheimer’s disease. Mol Neurodegener 9, 6. 10.1186/1750-1326-9-6.

39. Hammad, M.M., Dunn, H.A., and Ferguson, S.S.G. (2018). MAGI proteins can differentially regulate the signaling pathways of 5-HT2AR by enhancing receptor trafficking and PLC recruitment. Cellular Signalling 47, 109–121. 10.1016/j.cellsig.2018.03.016.

40. Pan-Vazquez, A., Wefelmeyer, W., Gonzalez Sabater, V., Neves, G., and Burrone, J. (2020). Activity-Dependent Plasticity of Axo-axonic Synapses at the Axon Initial Segment. Neuron 106, 265–276.e6. 10.1016/j.neuron.2020.01.037.

41. Burkarth, N., Kriebel, M., Kranz, E.U., and Volkmer, H. (2007). Neurofascin regulates the formation of gephyrin clusters and their subsequent translocation to the axon hillock of hippocampal neurons. Molecular and Cellular Neuroscience 36, 59–70. 10.1016/j.mcn.2007.06.001.

42. Kriebel, M., Metzger, J., Trinks, S., Chugh, D., Harvey, R.J., Harvey, K., and Volkmer, H. (2011). The Cell Adhesion Molecule Neurofascin Stabilizes Axo-axonic GABAergic Terminals at the Axon Initial Segment*. Journal of Biological Chemistry 286, 24385–24393. 10.1074/jbc.M110.212191.

43. Gao, Y., Hisey, E., Bradshaw, T.W.A., Erata, E., Brown, W.E., Courtland, J.L., Uezu, A., Xiang, Y., Diao, Y., and Soderling, S.H. (2019). Plug-and-Play Protein Modification Using Homology-Independent Universal Genome Engineering. Neuron 103, 583–597.e8. 10.1016/j.neuron.2019.05.047.

44. Willems, J., Jong, A.P.H. de, Scheefhals, N., Mertens, E., Catsburg, L.A.E., Poorthuis, R.B., Winter, F. de, Verhaagen, J., Meye, F.J., and MacGillavry, H.D. (2020). ORANGE: A CRISPR/Cas9-based genome editing toolbox for epitope tagging of endogenous proteins in neurons. PLOS Biology 18, e3000665. 10.1371/journal.pbio.3000665.

45. Droogers, W.J., Willems, J., MacGillavry, H.D., and Jong, A.P.H. de (2022). Duplex Labeling and Manipulation of Neuronal Proteins Using Sequential CRISPR/Cas9 Gene Editing. eNeuro 9. 10.1523/ENEURO.0056-22.2022.

46. Jamann, N., Dannehl, D., Lehmann, N., Wagener, R., Thielemann, C., Schultz, C., Staiger, J., Kole, M.H.P., and Engelhardt, M. (2021). Sensory input drives rapid homeostatic scaling of the axon initial segment in mouse barrel cortex. Nat Commun 12, 23. 10.1038/s41467-020-20232-x.

47. Galliano, E., Hahn, C., Browne, L.P., Villamayor, P.R., Tufo, C., Crespo, A., and Grubb, M.S. (2021). Brief Sensory Deprivation Triggers Cell Type-Specific Structural and Functional Plasticity in Olfactory Bulb Neurons. J. Neurosci. 41, 2135–2151. 10.1523/JNEUROSCI.1606-20.2020.

48. Kuba, H., Oichi, Y., and Ohmori, H. (2010). Presynaptic activity regulates Na+ channel distribution at the axon initial segment. Nature 465, 1075–1078. 10.1038/nature09087.

49. Greene, D.L., Kang, S., and Hoshi, N. (2017). XE991 and Linopirdine Are State-Dependent Inhibitors for Kv7/KCNQ Channels that Favor Activated Single Subunits. The Journal of Pharmacology and Experimental Therapeutics 362, 177–185. 10.1124/jpet.117.241679.

50. Tukker, A.M., Vrolijk, M.F., van Kleef, R.G.D.M., Sijm, D.T.H.M., and Westerink, R.H.S. (2023). Mixture effects of tetrodotoxin (TTX) and drugs targeting voltage-gated sodium channels on spontaneous neuronal activity *in vitro*. Toxicology Letters 373, 53–61. 10.1016/j.toxlet.2022.11.005.

51. Morris, R.G.M., Anderson, E., Lynch, G.S., and Baudry, M. (1986). Selective impairment of learning and blockade of long-term potentiation by an N-methyl-D-aspartate receptor antagonist, AP5. Nature 319, 774–776. 10.1038/319774a0.

52. Alford, S., and Grillner, S. (1990). CNQX and DNQX block non-NMDA synaptic transmission but not NMDA-evoked locomotion in lamprey spinal cord. Brain Research 506, 297–302. 10.1016/0006-8993(90)91266-J.

53. Kilman, V., Rossum, M.C.W. van, and Turrigiano, G.G. (2002). Activity Deprivation Reduces Miniature IPSC Amplitude by Decreasing the Number of Postsynaptic GABAA Receptors Clustered at Neocortical Synapses. J. Neurosci. 22, 1328–1337. 10.1523/JNEUROSCI.22-04-01328.2002.

54. Ibata, K., Sun, Q., and Turrigiano, G.G. (2008). Rapid Synaptic Scaling Induced by Changes in Postsynaptic Firing. Neuron 57, 819–826. 10.1016/j.neuron.2008.02.031.

55. Wefelmeyer, W., Cattaert, D., and Burrone, J. (2015). Activity-dependent mismatch between axo-axonic synapses and the axon initial segment controls neuronal output. Proceedings of the National Academy of Sciences 112, 9757–9762. 10.1073/pnas.1502902112.

56. Higgins, K.P., Naqib, V.A.B.A., Mayo, P., Lodder, B., Masuda, T., Amann, L., Prinz, M., and Kole, M.H.P. (2026). An organotypic neocortical slice culture for studying neuroglial interactions. Preprint at bioRxiv, 10.64898/2026.05.15.725074 10.64898/2026.05.15.725074.

57. Futai, K., Doty, C.D., Baek, B., Ryu, J., and Sheng, M. (2013). Specific Trans-Synaptic Interaction with Inhibitory Interneuronal Neurexin Underlies Differential Ability of Neuroligins to Induce Functional Inhibitory Synapses. J. Neurosci. 33, 3612–3623. 10.1523/JNEUROSCI.1811-12.2013.

58. Poulopoulos, A., Aramuni, G., Meyer, G., Soykan, T., Hoon, M., Papadopoulos, T., Zhang, M., Paarmann, I., Fuchs, C., Harvey, K., et al. (2009). Neuroligin 2 Drives Postsynaptic Assembly at Perisomatic Inhibitory Synapses through Gephyrin and Collybistin. Neuron 63, 628–642. 10.1016/j.neuron.2009.08.023.

59. Poulopoulos, A., Soykan, T., Tuffy, L.P., Hammer, M., Varoqueaux, F., and Brose, N. (2012). Homodimerization and isoform-specific heterodimerization of neuroligins. Biochem J 446, 321–330. 10.1042/BJ20120808.

60. Hirst, J., Borner, G.H.H., Harbour, M., and Robinson, M.S. (2005). The Aftiphilin/p200/γ-Synergin Complex. MBoC 16, 2554–2565. 10.1091/mbc.e04-12-1077.

61. Davidson, H.T., Xiao, J., Dai, R., and Bergson, C. (2009). Calcyon is necessary for activity-dependent AMPA receptor internalization and LTD in CA1 neurons of hippocampus. European Journal of Neuroscience 29, 42–54. 10.1111/j.1460-9568.2008.06563.x.

62. Yap, C.C., Digilio, L., McMahon, L., and Winckler, B. (2017). The endosomal neuronal proteins Nsg1/NEEP21 and Nsg2/P19 are itinerant, not resident proteins of dendritic endosomes. Sci Rep 7, 10481. 10.1038/s41598-017-07667-x.

63. Porat-Shliom, N., Milberg, O., Masedunskas, A., and Weigert, R. (2013). Multiple roles for the actin cytoskeleton during regulated exocytosis. Cell. Mol. Life Sci. 70, 2099–2121. 10.1007/s00018-012-1156-5.

64. Wernert, F., Moparthi, S.B., Pelletier, F., Lainé, J., Simons, E., Moulay, G., Rueda, F., Jullien, N., Benkhelifa-Ziyyat, S., Papandréou, M.-J., et al. (2024). The actin-spectrin submembrane scaffold restricts endocytosis along proximal axons. Science 385, eado2032. 10.1126/science.ado2032.

65. Wiesner, T., Parperis, C., Boroni-Rueda, F., Jullien, N., Mendes, A., Marie, L., Mezache, L., Papandréou, M.-J., Henriques, R., and Leterrier, C. (2025). Non-synaptic exocytosis along the axon shaft and its regulation by the submembrane periodic skeleton. Preprint at bioRxiv, 10.1101/2025.09.17.676728 10.1101/2025.09.17.676728.

66. Barry, J., Gu, Y., Jukkola, P., O’Neill, B., Gu, H., Mohler, P.J., Rajamani, K.T., and Gu, C. (2014). Ankyrin-G directly binds to kinesin-1 to transport voltage-gated Na+ channels into axons. Dev Cell 28, 117–131. 10.1016/j.devcel.2013.11.023.

67. Tai, Y., Gallo, N.B., Wang, M., Yu, J.-R., and Aelst, L.V. (2019). Axo-axonic Innervation of Neocortical Pyramidal Neurons by GABAergic Chandelier Cells Requires AnkyrinG-Associated L1CAM. Neuron 102, 358–372.e9. 10.1016/j.neuron.2019.02.009.

68. Ogawa, Y., Lim, B.C., George, S., Oses-Prieto, J.A., Rasband, J.M., Eshed-Eisenbach, Y., Hamdan, H., Nair, S., Boato, F., Peles, E., et al. (2023). Antibody-directed extracellular proximity biotinylation reveals that Contactin-1 regulates axo-axonic innervation of axon initial segments. Nat Commun 14, 6797. 10.1038/s41467-023-42273-8.

69. Jung, K., Choi, Y., and Kwon, H.-B. (2022). Cortical control of chandelier cells in neural codes. Front. Cell. Neurosci. 16. 10.3389/fncel.2022.992409.

70. Seignette, K., Jamann, N., Papale, P., Terra, H., Porneso, R.O., de Kraker, L., van der Togt, C., van der Aa, M., Neering, P., Ruimschotel, E., et al. (2024). Experience shapes chandelier cell function and structure in the visual cortex. eLife 12, RP91153. 10.7554/eLife.91153.

71. Jung, K., Chang, M., Steinecke, A., Burke, B., Choi, Y., Oisi, Y., Fitzpatrick, D., Taniguchi, H., and Kwon, H.-B. (2023). An adaptive behavioral control motif mediated by cortical axo-axonic inhibition. Nat Neurosci 26, 1379–1393. 10.1038/s41593-023-01380-x.

72. Zhao, R., Ren, B., Xiao, Y., Tian, J., Zou, Y., Wei, J., Qi, Y., Hu, A., Xie, X., Huang, Z.J., et al. (2024). Axo-axonic synaptic input drives homeostatic plasticity by tuning the axon initial segment structurally and functionally. Science Advances 10, eadk4331. 10.1126/sciadv.adk4331.

73. DeFelipe, J., Sola, R.G., Marco, P., Rio, M. del R. del, Pulido, P., and Cajal, S.R. y (1993). Selective Changes in the Microorganization of the Human Epileptogenic Neocortex Revealed by Parvalbumin Immunoreactivity. Cereb Cortex 3, 39–48. 10.1093/cercor/3.1.39.

74. Woo, T.-U., Whitehead, R.E., Melchitzky, D.S., and Lewis, D.A. (1998). A subclass of prefrontal γ-aminobutyric acid axon terminals are selectively altered in schizophrenia. Proceedings of the National Academy of Sciences 95, 5341–5346. 10.1073/pnas.95.9.5341.

75. Ariza, J., Rogers, H., Hashemi, E., Noctor, S.C., and Martínez-Cerdeño, V. (2018). The Number of Chandelier and Basket Cells Are Differentially Decreased in Prefrontal Cortex in Autism. Cereb Cortex 28, 411–420. 10.1093/cercor/bhw349.

76. Gallo, N.B., Paul, A., and Van Aelst, L. (2020). Shedding Light on Chandelier Cell Development, Connectivity, and Contribution to Neural Disorders. Trends in Neurosciences 43, 565–580. 10.1016/j.tins.2020.05.003.

77. Vivien, J., El Azraoui, A., Lheraux, C., Lanore, F., Aouizerate, B., Herry, C., Humeau, Y., and Bienvenu, T.C.M. (2023). Axo-axonic cells in neuropsychiatric disorders: a systematic review. Front. Cell. Neurosci. 17. 10.3389/fncel.2023.1212202.

78. Oguro, K., Shimazaki, K., Yokota, H., Onuki, Y., Murashima, Y., Kawai, K., and Muramatsu, S. (2022). Global brain delivery of neuroligin 2 gene ameliorates seizures in a mouse model of epilepsy. The Journal of Gene Medicine 24, e3402. 10.1002/jgm.3402.

79. Parente, D.J., Garriga, C., Baskin, B., Douglas, G., Cho, M.T., Araujo, G.C., and Shinawi, M. (2017). Neuroligin 2 nonsense variant associated with anxiety, autism, intellectual disability, hyperphagia, and obesity. American Journal of Medical Genetics Part A 173, 213–216. 10.1002/ajmg.a.37977.

80. Trobiani, L., Meringolo, M., Diamanti, T., Bourne, Y., Marchot, P., Martella, G., Dini, L., Pisani, A., De Jaco, A., and Bonsi, P. (2020). The neuroligins and the synaptic pathway in Autism Spectrum Disorder. Neuroscience & Biobehavioral Reviews 119, 37–51. 10.1016/j.neubiorev.2020.09.017.

81. van Beuningen, van Beuningen, S.F.B., Lena Will, Will, L., Martin Harterink, Harterink, M., Anaël Chazeau, Chazeau, A., Eljo Y. van Battum, van Battum, E.Y., et al. (2015). TRIM46 Controls Neuronal Polarity and Axon Specification by Driving the Formation of Parallel Microtubule Arrays. Neuron 88, 1208–1226. 10.1016/j.neuron.2015.11.012.

82. Kim, E., Niethammer, M., Rothschild, A., Nung Jan, Y., and Sheng, M. (1995). Clustering of Shaker-type K+ channels by interaction with a family of membrane-associated guanylate kinases. Nature 378, 85–88. 10.1038/378085a0.

83. Lois, C., Hong, E.J., Pease, S., Brown, E.J., and Baltimore, D. (2002). Germline Transmission and Tissue-Specific Expression of Transgenes Delivered by Lentiviral Vectors. Science 295, 868–872. 10.1126/science.1067081.

84. Stewart, S.A., Dykxhoorn, D.M., Palliser, D., Mizuno, H., Yu, E.Y., An, D.S., Sabatini, D.M., Chen, I.S.Y., Hahn, W.C., Sharp, P.A., et al. (2003). Lentivirus-delivered stable gene silencing by RNAi in primary cells. RNA 9, 493–501. 10.1261/rna.2192803.

85. Weigel, A.V., Chang, C.-L., Shtengel, G., Xu, C.S., Hoffman, D.P., Freeman, M., Iyer, N., Aaron, J., Khuon, S., Bogovic, J., et al. (2021). ER-to-Golgi protein delivery through an interwoven, tubular network extending from ER. Cell 184, 2412–2429.e16. 10.1016/j.cell.2021.03.035.

86. Chih, B., Gollan, L., and Scheiffele, P. (2006). Alternative Splicing Controls Selective Trans-Synaptic Interactions of the Neuroligin-Neurexin Complex. Neuron 51, 171–178. 10.1016/j.neuron.2006.06.005.

87. Farías, G.G., Guardia, C.M., De Pace, R., Britt, D.J., and Bonifacino, J.S. (2017). BORC/kinesin-1 ensemble drives polarized transport of lysosomes into the axon. Proceedings of the National Academy of Sciences 114, E2955–E2964. 10.1073/pnas.1616363114.

88. Stirling, D.R., Swain-Bowden, M.J., Lucas, A.M., Carpenter, A.E., Cimini, B.A., and Goodman, A. (2021). CellProfiler 4: improvements in speed, utility and usability. BMC Bioinformatics 22, 433. 10.1186/s12859-021-04344-9.

89. Farías, G.G., Fréal, A., Tortosa, E., Stucchi, R., Pan, X., Portegies, S., Will, L., Altelaar, M., and Hoogenraad, C.C. (2019). Feedback-Driven Mechanisms between Microtubules and the Endoplasmic Reticulum Instruct Neuronal Polarity. Neuron 102, 184–201.e8. 10.1016/j.neuron.2019.01.030.

90. Concordet, J.-P., and Haeussler, M. (2018). CRISPOR: intuitive guide selection for CRISPR/Cas9 genome editing experiments and screens. Nucleic Acids Res 46, W242–W245. 10.1093/nar/gky354.

91. Schwartz, M.L., and Jorgensen, E.M. (2016). SapTrap, a Toolkit for High-Throughput CRISPR/Cas9 Gene Modification in Caenorhabditis elegans. Genetics 202, 1277–1288. 10.1534/genetics.115.184275.

92. Cox, J., and Mann, M. (2008). MaxQuant enables high peptide identification rates, individualized p.p.b.-range mass accuracies and proteome-wide protein quantification. Nat Biotechnol 26, 1367–1372. 10.1038/nbt.1511.

93. Koopmans, F., van Nierop, P., Andres-Alonso, M., Byrnes, A., Cijsouw, T., Coba, M.P., Cornelisse, L.N., Farrell, R.J., Goldschmidt, H.L., Howrigan, D.P., et al. (2019). SynGO: An Evidence-Based, Expert-Curated Knowledge Base for the Synapse. Neuron 103, 217–234.e4. 10.1016/j.neuron.2019.05.002.

94. Arshadi, C., Günther, U., Eddison, M., Harrington, K.I.S., and Ferreira, T.A. (2021). SNT: a unifying toolbox for quantification of neuronal anatomy. Nat Methods 18, 374–377. 10.1038/s41592-021-01105-7.

95. Ollion, J., Cochennec, J., Loll, F., Escudé, C., and Boudier, T. (2013). TANGO: a generic tool for high-throughput 3D image analysis for studying nuclear organization. Bioinformatics 29, 1840–1841. 10.1093/bioinformatics/btt276.

